# Temperature-dependent effects of house fly proto-Y chromosomes on gene expression

**DOI:** 10.1101/2020.08.05.238378

**Authors:** Kiran Adhikari, Jae Hak Son, Anna H. Rensink, Jaweria Jaweria, Daniel Bopp, Leo W. Beukeboom, Richard P. Meisel

**Affiliations:** Department of Biology and Biochemistry, University of Houston, Houston, TX USA 77204-5001; Groningen Institute for Evolutionary Life Sciences, University of Groningen, 9700 CC Groningen, The Netherlands; Institute of Molecular Life Sciences, University of Zurich, Switzerland CH-8057; Department of Epidemiology of Microbial Diseases, Yale University School of Public Health, New Haven, CT 06520

## Abstract

Sex determination, the developmental process by which sexually dimorphic phenotypes are established, evolves fast. Species with polygenic sex determination, in which master regulatory genes are found on multiple different proto-sex chromosomes, are informative models to study the evolution of sex determination. House flies are such a model system, with male determining loci possible on all six chromosomes and a female-determiner on one of the chromosomes as well. The two most common male-determining proto-Y chromosomes form latitudinal clines on multiple continents, suggesting that temperature variation is an important selection pressure responsible for maintaining polygenic sex determination in this species. To identify candidate genes that may be under selection, we used RNA-seq to test for temperature-dependent effects of the proto-Y chromosomes on gene expression in adult house flies. We find no evidence for ecologically meaningful temperature-dependent expression of sex determining genes between male genotypes, but we were likely not sampling an appropriate developmental time-point to identify such effect. In contrast, we identified many other genes whose expression depends on the interaction between proto-Y chromosome genotype and temperature, including genes that encode proteins involved in reproduction, metabolism, lifespan, stress response, and immunity. Notably, genes with genotype-by-temperature interactions on expression are not enriched on the proto-sex chromosomes. Moreover, there is no evidence that temperature-dependent expression is driven by chromosome-wide expression divergence between the proto-Y and proto-X alleles. Therefore, if temperature-dependent gene expression is responsible for differences in phenotypes and fitness of proto-Y genotypes across house fly populations, these effects are driven by a small number of temperature-dependent alleles on the proto-Y chromosomes that may in turn affect the expression of genes on other chromosomes.

## Introduction

Sex determination establishes sexually dimorphic developmental pathways, either based on genetic differences between males and females or environmental cues (Beukeboom & Perrin 2014). In species with genotypic sex determination, a single master regulatory locus (e.g., *SRY* on the human Y chromosome) is often enough to initiate whether the embryo develops into a male or female (Sinclair *et al*. 1990; Goodfellow & Lovell-Badge 1993). However, in polygenic sex determination systems, multiple master sex determining loci segregate independently, often on different chromosomes (Moore & Roberts 2013). Most population genetics models predict that polygenic sex determination will be an evolutionary intermediate between different monogenic sex determination systems, and the factors responsible for maintaining polygenic sex determination as a stable polymorphism are poorly understood (Rice 1986; van Doorn 2014). Models that do allow for the stable maintenance of polygenic sex determination require opposing (sexually antagonistic) fitness effects of sex chromosomes in males and females (Orzack *et al*. 1980; van Doorn & Kirkpatrick 2007, 2010). Understanding how other selection pressures can maintain polygenic sex determination would provide valuable insight into the factors that drive the evolution of sex determination.

House fly (*Musca domestica*) is a well suited model for studying polygenic sex determination because multiple male and female determining loci segregate on different chromosomes in natural populations (Hamm *et al*. 2015). Male sex in house fly is initiated by the gene *Musca domestica male determiner, Mdmd* (Sharma *et al*. 2017). *Mdmd* arose via the recent duplication of the ubiquitous splicing factor *nucampholin* (*Md-ncm*) after the divergence of house fly from its close relative *Stomoxys calcitrans*. *Mdmd* promotes male development by causing the house fly ortholog of *transformer* (*Md-tra*) to be spliced into non-functional isoforms with premature stop codons (Hediger *et al*. 2010). The lack of functional Md-Tra protein leads to male-specific splicing of *doublesex* (*Md-dsx*) and *fruitless* (*Md-fru*), the two known downstream targets of *Md-tra* (Hediger *et al*. 2004; Meier *et al*. 2013). In the absence of *Mdmd*, *Md-tra* is spliced into a functional transcript that is translated into a protein that promotes female specific splicing of *Md-dsx* and inhibits splicing of the male isoform of *Md-fru*.

*Mdmd* can be found on multiple different chromosomes in house fly (Sharma *et al*. 2017), and it is most commonly found on the third (III^M^) and Y (Y^M^) chromosomes (Hamm *et al*. 2015). While Y^M^ is conventionally referred to as the Y chromosome, both III^M^ and Y^M^ are young proto-Y chromosomes that are minimally differentiated from their homologous proto-X chromosomes (Meisel *et al*. 2017; Son & Meisel 2021). The proto-Y chromosomes are clinally distributed—with III^M^ most common at southern latitudes and Y^M^ most common at northern latitudes—across multiple continents (Hiroyoshi 1964; Mcdonald *et al*. 1975; Denholm *et al*. 1986; Hamm *et al*. 2005). The frequencies of III^M^ and Y^M^ in natural populations have remained stable for decades (Kozielska *et al*. 2008; Meisel *et al*. 2016). This clinal distribution of III^M^ and Y^M^, along with their stable frequencies across populations, suggests that natural selection maintains the polymorphism.

A female determining allele of *Md-tra* (*Md-tra^D^*) is also found in some house fly populations (McDonald *et al*. 1978; Hediger *et al*. 2010). *Md-tra^D^* can initiate female development in embryos with at least three *Mdmd* chromosomes (Schmidt *et al*. 1997; Hediger *et al*. 1998). In some populations, both Y^M^ and III^M^ can be found, with some males carrying one copy of two different proto-Y chromosomes or homozygous for a proto-Y (e.g., Hamm & Scott 2008, 2009). *Md-tra^D^* is most common in populations with a high frequency of these “multi-Y” males, which results in a sex ratio with an equal number of males and females (Meisel *et al*. 2016).

The natural distribution of III^M^ and Y^M^ hints at a possible genotype-by-temperature (G×T) interaction that could explain the stable maintenance of Y^M^-III^M^ clines. Temperature is not the only selection pressure that could vary along the clines, but seasonality in temperature is the best predictor of the frequencies of the proto-Y chromosomes across populations (Feldmeyer *et al*. 2008). There are at least two non exclusive ways in which temperature-dependent selection pressures could maintain the III^M^-Y^M^ polymorphism. First, alleles on the III^M^ and Y^M^ chromosomes (other than the *Mdmd* locus) could have temperature-dependent fitness effects. In this scenario, the III^M^-Y^M^ clines would be maintained in a similar way to how temperature variation maintains opposing clines of heat and cold tolerance in *Drosophila melanogaster* between tropical and temperate regions (Hoffmann *et al*. 2002). Second, it is possible that the *Mdmd* copies on the III^M^ and Y^M^ chromosomes differ in their temperature dependent activities, such that *Mdmd* on the III^M^ chromosome increases male fitness at warm temperatures and *Mdmd* on the Y^M^ chromosome increases fitness at colder temperatures. This is analagous to how, in some fish and reptile species, temperature can drive sex determination and override the outcomes of genotypic sex determining systems (Shine *et al*. 2002; Quinn *et al*. 2007; Radder *et al*. 2008; Holleley *et al*. 2015).

We investigated if temperature-dependent phenotypic effects of III^M^ and Y^M^ could be driven by differential gene expression in males across temperatures. We selected gene expression as a phenotypic read-out of G×T interactions because temperature-dependent differences in gene expression are well documented in clinally distributed genetic variation (Levine *et al*. 2011; Zhao *et al*. 2015). Specifically, we evaluated how G×T interactions affect gene expression in male house flies carrying either a III^M^ or Y^M^ chromosome. We used RNA-seq to study gene expression in two nearly isogenic lines of house flies, differing only by their proto-Y chromosome, reared at two developmental temperatures. This allowed us to assess the effects of the entire III^M^ and Y^M^ chromosomes. We also used quantitative reverse transcription PCR (qRT-PCR) to investigate the temperature-dependent expression of *Mdmd*.

## Materials & Methods

### qRT-PCR samples and analysis

We used qRT-PCR to measure the expression of *Mdmd* and its paralog *Md-ncm* in two Y^M^ strains and two III^M^ strains. The strains were grouped into two pairs, with one Y^M^ strain and one III^M^ strain per pair. In the first pair, we used the Y^M^ strain IsoCS and the III^M^ strain CSkab (both from North America). IsoCS and CSkab share a common genetic background of the Cornell susceptible (CS) strain (Scott *et al*. 1996). IsoCS was previously created by crossing a Y^M^ chromosome from Maine onto the CS background (Hamm *et al*. 2009). We created CSkab by backcrossing the III^M^ chromosome from the KS8S3 strain collected in Florida (Kaufman *et al*. 2010) onto the CS background, using an approach described previously (Son *et al*. 2019). In the second pair, we used two European strains: the Y^M^ strain GK-1 from Gerkesklooster (Netherlands) and the III^M^ strain SPA3 from near Girona (Spain). GK-1 and SPA3 were maintained in the lab, each as inbred populations, for approximately 40 and 50 generations, respectively.

We raised all strains at 18°C and 27°C for two generations with 12:12-h light:dark photoperiods. Adult males and females for each GxT combination were housed in cages with *ad libitum* containers of 1:1 combinations of sugar and non-fat dry milk and *ad libitum* containers of water. Females were provided with a standard medium of wheat bran, calf manna, wood chips, yeast, and water in which to lay eggs for 12-24 hrs (Hamm *et al*. 2009). The resulting larvae were maintained in the same media within 32 oz containers. Adult females did not lay a sufficient number of eggs at 18°C, so the adults from the 18°C colonies were transferred to 22°C for egg laying for 1-2 days. The eggs collected at 22°C were then moved back to 18°C for larval development, pupation, and emergence as adults. We maintained the colonies at these temperatures for two generations. Collecting flies after two generations ensured at least one full egg-to-adult generation at the appropriate temperature.

For qRT-PCR experiments involving the North American IsoCS and CSkab strains, abdomen samples were dissected from 5 day old adult males after being anesthetized with CO_2_. For qRT-PCR assessments on the European GK and SPA3 strains, full body samples were collected from 5 day old adult males after being anesthetized with CO_2_. Tissue samples from 5-7 males were pooled in each of three biological replicates for each genotype (Y^M^ and III^M^) by temperature (18°C and 27°C) combination. The collected tissues were homogenized in TRIzol reagent (Life Technologies) using a motorized grinder in a 1.5 ml microcentrifuge tube. For the North American strains, the Direct-zol RNA MiniPrep kit (Zymo Research) was used to extract RNA from the homogenized samples. The isolated RNA was reverse transcribed into cDNA with MLV RT (Promega), following the manufacturer’s protocol. For the European strains, the RNA phase following centrifugation with TRIzol reagent was separated using chloroform and precipitated by using isopropanol and ethanol. The isolated RNA was reverse transcribed into cDNA using RevertAid H minus 1st strand kit (Fermentas #K1632) according to the manufacturer’s protocol.

We conducted qRT-PCR of cDNA from the male flies. We used qRT-PCR primers (Supplementary Table 1) to uniquely amplify *Mdmd* and *Md-ncm* without amplifying the other paralog (Sharma *et al*. 2017). Primers were additionally used to amplify cDNA from a transcript (LOC101888902) that is not differentially expressed between Y^M^ and III^M^ males as an internal control for cDNA content in each biological replicate (Meisel *et al*. 2015). The IsoCS and CSkab samples were assayed on a StepOnePlus machine using PowerUp SYBR Green Master Mix (Applied Biosystems). The GK and SPA3 samples were assayed on a Applied Biosystems qPCR cycler 7300 machine using Quanta perfecta SYBR Green Fastmix (Quanta bio). We measured the abundance of PCR products from each primer pair in three technical replicates of three biological replicates for each G×T combination. With the same primer pairs, we also measured the expression of serial dilutions (1/1, 1/5, 1/25, 1/125, and 1/625) of cDNA from independent biological collections of house flies. Samples were interspersed across 96-well microtiter plates to minimize batch effects.

We constructed standard curves for each primer pair by calculating the linear relationship between CT values and log_10_(concentration) from the serial dilutions using the lm() function in the R statistical programming package (R Core Team 2019). We then used the equations of the standard curves to calculate the concentration of transcripts (i.e., cDNA) from *Mdmd* and *Md-ncm* in each technical replicate. We next determined a normalized expression level of each technical replicate by dividing the concentration of the technical replicate by the mean concentration of the control transcript (LOC101888902) across the three technical replicates from the same biological replicate.

We used an analysis of variance (ANOVA) approach to test for the effect of genotype (Y^M^ vs III^M^), developmental temperature (18°C vs 27°C), and the interaction of genotype and temperature on the expression of each transcript. To those ends, we used the lmer() function in the lme4 package (Bates *et al*. 2015) in R to model the effect of genotype (*G*), temperature (*T*), and the interaction term as fixed effect factors, as well as biological replicate (*r*) as a random effect, on expression level (*E*):

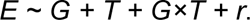

We then compared the fit of that full model to a model without the interaction term (*E ∼ G + T + r*) using the anova() function in R. If the full model fits significantly better, that is evidence that there is a significant G*×*T interaction on the expression of the transcript.

### RNA-seq samples

We used RNA-seq to measure gene expression in the Y^M^ strain IsoCS and a III^M^ strain known as CSrab. IsoCS (described above) and CSrab have different proto-Y chromosomes on the shared CS genetic background (Scott *et al*. 1996). We created CSrab by backcrossing the III^M^ chromosome of a spinosad-resistant strain, rspin (Shono & Scott 2003), onto the CS background, using the same approach as we used to create CSkab, described elsewhere (Son *et al*. 2019). These strains are normally raised at 25°C, but were raised at different temperatures (18°C or 29°C) in our experiment in order to determine the effect of genotype and temperature on gene expression.

Colonies of both strains were reared at 18°C and 29°C for two generations with at least one full egg-to-adult generation, as described above. We therefore had four combinations of genotype (Y^M^ and III^M^) and temperature (18°C and 29°C). We controlled for the adult density using 35 adult males and 35 adult females for each G×T combination. We also controlled for larval density with 100 larvae per 32 oz container. Third generation males obtained from second generation females were collected and reared separately from the females at their respective developmental temperatures for 1–8 days before RNA extraction.

For the RNA-seq experiments, head and testis samples from 1–8 day old males were dissected in 1% PBS solution after being anesthetized with CO_2_. We dissected testes from 15–20 house flies per each of three replicates of each G×T combination. Similarly, 5 heads were dissected for each of three biological replicates for each G×T combination. The collected tissues were homogenized in TRIzol reagent (Life Technologies) using a motorized grinder in a 1.5 mL microcentrifuge tube. The Direct-zol RNA MiniPrep kit (Zymo Research) was used to extract RNA from the homogenized samples. RNA-seq library preparation was carried out using the TruSeq Stranded mRNA Kit (Illumina). Qualities of these libraries were assessed using a 2100 Bioanalyzer (Agilent Technologies, Inc.). Libraries were then sequenced with 75 bp single-end reads on high output runs of an Illumina NextSeq 500 at the University of Houston Seq-N-Edit Core. All testis samples (i.e., all replicates of each G×T combination) were sequenced together in a single run, and all head samples were sequenced together on a separate run. All RNA-seq data are available in the NCBI Gene Expression Omnibus under accession GSE136188 (BioProject PRJNA561541, SRA accession SRP219410).

### RNA-seq data analysis

RNA-seq reads were aligned to the annotated house fly reference genome Musca_domestica-2.0.2 (Scott *et al*. 2014) using HISAT2 (Kim *et al*. 2015) with the default settings of a maximum mismatch penalty of 6 and minimum penalty of 2, and a soft-clip penalty of maximum 2 and minimum 1 (Supplementary Tables 2 and 3). We next used SAMtools (Li *et al*. 2009) to sort the aligned reads. The sorted reads were assigned to annotated genes (*M. domestica* Annotation Release 102) using htseq-count in HTSeq (Anders *et al*. 2015). We only included uniquely mapped reads, and we excluded reads with ambiguous mapping and reads with a mapping quality of less than 10.

We analyzed the exon-level expression of the sex determining genes *Md-tra* (LOC101888218) and *Md-dsx* (LOC101895413) for each G×T combination. To do so, we first determined the read coverage across *Md-tra* and *Md-dsx* transcripts using the ‘mpileup’ function in SAMtools (Li et al., 2009). We then calculated normalized read depth (*D_ijk_*) at each site *i* within each gene in library *j* for each G×T combination *k* by dividing the number of reads mapped to a site (*r_ijk_*) into the total number of reads mapped in that library (*R_jk_*), and we multiplied that value by one million:

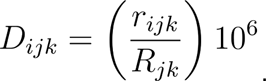

For each site within each gene, we then calculated the average *D_ijk_* across all three l ibraries for each G×T combination 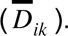

We also used the DESeq2 package in R (Love *et al*. 2014) to analyze differential expression of all annotated genes between all G×T combinations. To do so, we used a linear model that included genotype (Y^M^ or III^M^), developmental temperature, and their interaction term to predict gene expression levels:

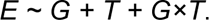

Genes for which the interaction term has a false discovery rate (FDR) corrected *P*-value (Benjamini & Hochberg 1995) of less than 0.05 were considered to be differentially expressed as a result of the G×T interaction. The same FDR corrected cutoff was used to test for genes that are differentially expressed according to genotype or temperature, by testing for the effect of *G* or *T* using results analyzed with the full model. For principal component analysis (PCA), hierarchical clustering, and non-metric multidimensional scaling (NMDS), we analyzed regularized log transformed count data generated by the rlog() function in DESeq2. NMDS was carried out using metaMDS() function from the vegan package in R with the autotransform = FALSE option (Oksanen *et al*. 2019).

We performed a gene ontology (GO) analysis to test for enrichment of functional classes amongst differentially expressed genes. To assign GO terms to house fly genes, we first used BLASTX to search house fly transcripts against a database of all *D. melanogaster* proteins (Gish & States 1993). We took this approach because GO assignments are missing for most house fly genes. The top hit for each house fly gene obtained from BLASTX was used to assign a FlyBase ID to each house fly transcript. These *D. melanogaster* homologs were then used in DAVID 6.8 (Huang *et al*. 2009a; b) to identify GO terms that are significantly enriched amongst differentially expressed genes (FDR corrected *P* < 0.05).

### Allele-specific expression analysis

We tested for differential expression of third chromosome genes between the allele on the III^M^ chromosome and the allele on the standard (non-*Mdmd*) third chromosome in III^M^ males. To do so, we followed the Genome Analysis Toolkit (GATK) best practices workflow for single nucleotide polymorphism (SNP) and insertion/deletion (indel) calling to identify sequence variants in our RNA-seq data (McKenna *et al*. 2010; Meisel *et al*. 2017). We first used STAR (Dobin *et al*. 2013) to align reads from the 12 testis libraries and 12 head libraries to the house fly reference genome (Musca_domestica-2.0.2). We then used the splice junction information from the first alignment to create a new index that was used to perform a second alignment. Using *de novo* transcripts identified with STAR serves to reduce read-mapping biases associated with an incomplete transcript annotation. After adding read group information to the SAM file thus generated, we marked duplicates. We next used SplitNCigarReads to reassign mapping qualities to 60 with the ReassignOneMappingQuality read filter for alignments with a mapping quality of 255. We used RealignerTargetCreator to identify and IndelRealigner to realign the indels. We used BaseRecalibrator and variant calls from a previous RNA-seq analysis (Meisel *et al*. 2017) to recalibrate the realigned reads. The realigned reads were then used for variant calling with HaplotypeCaller with emission and calling thresholds of 20. We filtered the variants obtained using VariantFiltration with a cluster window size of 35 bp, cluster size of 3 SNPs, FS > 30, and QD < 2. This filtering was applied because there may be preferential mapping of reads containing SNPs found in the reference genome relative to reads with alternative SNPs (Stevenson *et al*. 2013; Zimmer *et al*. 2016). By excluding SNPs found in clusters of at least 3 in a 35 bp window from our analysis, we can greatly reduce read-mapping biases from our estimates of allele-specific expression (Son & Meisel 2021).

We then used all the generated gvcf files to carry out joint genotyping using GenotypeGVCFs. We performed separate joint genotyping for testis and head libraries. The variants from Joint Genotyping were then filtered using VariantFiltration with FS > 30 and QD < 2. We used the vcfR package in R (Knaus & Grünwald 2017) to extract information from vcf files obtained from joint genotyping. For downstream analysis, we only kept SNPs (i.e., variants where the reference and alternate allele are 1 bp), and excluded small indels.

To test for allele-specific expression, we first assigned sequence variants to the III^M^ and standard third (III) chromosomes. This was only done for sites that were heterozygous in III^M^ males and homozygous in Y^M^ males (all other variable sites on the third chromosome were discarded) because these are the only alleles we can assign to either the III^M^ or III chromosome. This is because Y^M^ males are homozygous for the III chromosome (X/Y^M^; III/III), and III^M^ males are heterozygous (X/X; III^M^/III). For every variable site, we assigned the allele shared by both III^M^ and Y^M^ males to the III chromosome, and the allele unique to III^M^ males to the III^M^ chromosome. We calculated the sum of read depth for each allele across all three sequencing libraries (i.e., replicates) of each G×T combination. For each gene, we calculated the average normalized read depth across all variable sites within the gene separately for the III^M^ and III alleles at each temperature. To compare the expression of the III^M^ and III alleles, we calculated the difference in sequencing coverage between III^M^ and III alleles at each site for each temperature separately. We calculated the average difference in expression of III^M^ and III alleles in each gene at each temperature *k*, *d_k_*, as follows:

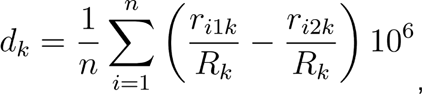

where *r_i_*_1*k*_ is the expression of the III^M^ allele at site *i* (out of *n* total polymorphic sites) andtemperature *k* (either 18°C or 29°C), *r_i_*_2*k*_ is the expression of the III allele at site *i* and temperature *k*, and *R_k_* is the total number of mapped reads in III^M^ males at temperature *k* . We then calculated standard error of *d_k_* across all sites for each gene at each temperature.

## Results

### Genotype and temperature affect genome-wide gene expression profiles

We used RNA-seq to test for the effects of genotype and developmental temperature on gene expression in heads and testes of Y^M^ and III^M^ house flies raised at 18°C and 29°C. The purpose of raising the strains at two different temperatures is to expose G×T effects of Y^M^ and III^M^ alleles sampled from natural populations (i.e., genotype-dependent plasticity across temperatures), not to evolve adaptations to each temperature. We first used PCA, NMDS, and hierarchical clustering to assess the similarities of the overall gene expression profiles of each of three replicates of each G×T combination in head and testis separately.

The PCA of the head RNA-seq data (using all 16,540 expressed genes) provides some evidence for an effect of genotype on gene expression. The first principal component (PC1) of head gene expression explains 34% of the variance in expression, and the second (PC2) explains 23% of the variation (Figure 1A). However, there is no clear grouping by genotype or developmental temperature, which can be best explained by an age-effect in our samples. One biological replicate of III^M^ heads at each temperature came from older males (4–8 days old, as opposed to the other samples which were 1–3 days old). The two older samples had head expression profiles that clustered separately from the remaining samples in our PCA (Figure 1A). Excluding the two older samples, we found a clear grouping by genotype along PC2, which explains 28% of the variance in head gene expression (Figure 1B). Because of the effect of age on head gene expression, we describe results both including and excluding the two older samples in the remainder of the analyses we present.

**Figure 1.**
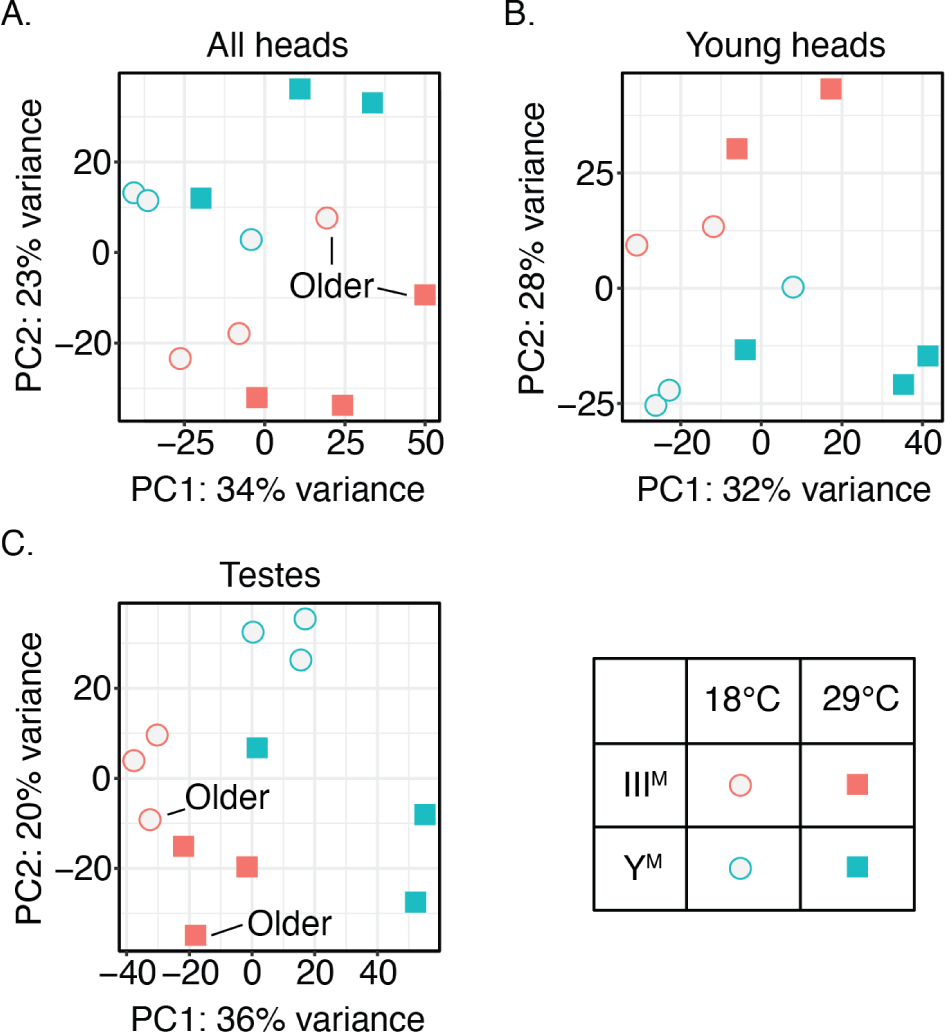
Effect of genotype and temperature on genome-wide gene expression in house flies. Graphs show the first two principal components (PC) explaining gene expression levels in all male heads (A), heads of young males only (B) and testes (C) samples. Each data point represents a biological replicate, with PC coordinates determined using regularized log transformed read counts.

In testis, PC1 explains 36% of the variance in expression, and it separates III^M^ males at 18°C from Y^M^ males at 29°C (Figure 1C). III^M^ is found at southern, warmer temperatures, whereas Y^M^ is found at northern, colder temperatures. PC1 for testis expression therefore separates the two genotypes at the temperatures that are opposite from their geographic distribution (i.e., Y^M^ occurs at relatively low temperature and III^M^ at high temperature). PC2 explains 20% of the variation in testis expression and separates III^M^ at 29°C from Y^M^ at 18°C (Figure 1C). Therefore, PC2 separates the two genotypes at temperatures that are consistent with their geographic distribution. We did not observe a meaningful effect of age on gene expression in testis (Figure 1C), and we thus did not repeat the analysis excluding the older testis samples.

We performed the following analyses to evaluate the robustness of our PCA results. First, we carried out PCA by considering only the 500 most variable genes in head and testis and observed the same patterns as those described above (Supplementary Figure S1). We additionally carried out PCA for genes on each chromosome, and the results for each chromosome were consistent with those across all chromosomes (Supplementary Figures S2, S3, and S4). Notably, there is very strong differentiation of III^M^ and Y^M^ males when we consider the testis expression of X chromosome and third chromosome genes (Supplementary Figure S4). This can be explained by the fact that the two genotypes only differ in these chromosomes, and share the same genetic background for the remaining chromosomes. We also carried out NMDS and hierarchical clustering of the RNA-seq data. We observed a grouping by genotype in the NMDS for head samples, and grouping by genotype and temperature in the testis samples (Supplementary Figure S5). In the hierarchical clustering, we did not observe grouping by genotype or temperature in head samples while including or excluding the older samples (Supplementary Figure S6). For testis gene expression, we found some evidence for clustering first by genotype and then by temperature (Supplemental Figure S6), similar to the PCA. However, the concordance between clusters and G×T combinations is not perfect.

### Genotype and temperature affect the expression of individual genes

To further test for genotype- and temperature-dependent gene expression, we next identified differentially expressed genes in two types of pairwise comparisons: i) between genotypes at one developmental temperature (either at 18°C or 29°C), and ii) within a genotype across the two developmental temperatures. Comparing between genotypes, we found 900 genes that are differentially expressed between Y^M^ and III^M^ heads at 18°C, and there were 1378 genes differentially expressed between Y^M^ and III^M^ heads at 29°C (Supplementary Table 4, Supplementary Figure S7). Excluding the two older samples, we found 786 genes that are differentially expressed between Y^M^ and III^M^ heads at 18°C, and 1748 genes differentially expressed between Y^M^ and III^M^ heads at 29°C (Supplementary Table 5, Supplementary Figure S7). The increase in differentially expressed genes at 29°C when the older samples are excluded can be explained by reduced variation within the III^M^ male samples, which should increase our power to detect differences between III^M^ and Y^M^ males. The number of differentially expressed genes is higher in testis than head: 2413 genes at 18°C and 2199 genes at 29°C are significantly differentially expressed between Y^M^ and III^M^ testes (Supplementary Table 6, Supplementary Figure S7). This is consistent with previous work that identified more genes differentially expressed between Y^M^ and III^M^ males in testis than head (Meisel *et al*. 2015).

In both head and testis, there is an excess of genes on the third chromosome that are significantly differentially expressed between genotypes at both 18°C and 29°C (Figure 2A), regardless of whether the older samples are excluded (Supplementary Figure S8). This is consistent with different third chromosome genotypes between strains, and it is suggestive of a *cis* effect on gene expression levels (Meisel *et al*. 2015; Son *et al*. 2019). The excess differential expression of chromosome III genes is also consistent with the signal that chromosome III gene expression provides to differentiating III^M^ and Y^M^ males (Supplementary Figure S4). The observed proportion of differentially expressed X-linked genes also appears to deviate from the expectation based on the genome-wide average (Figure 2A), but it is not significant because of low power caused by the small number (<100) of genes on the house fly X chromosome (Meisel & Scott 2018).

**Figure 2.**
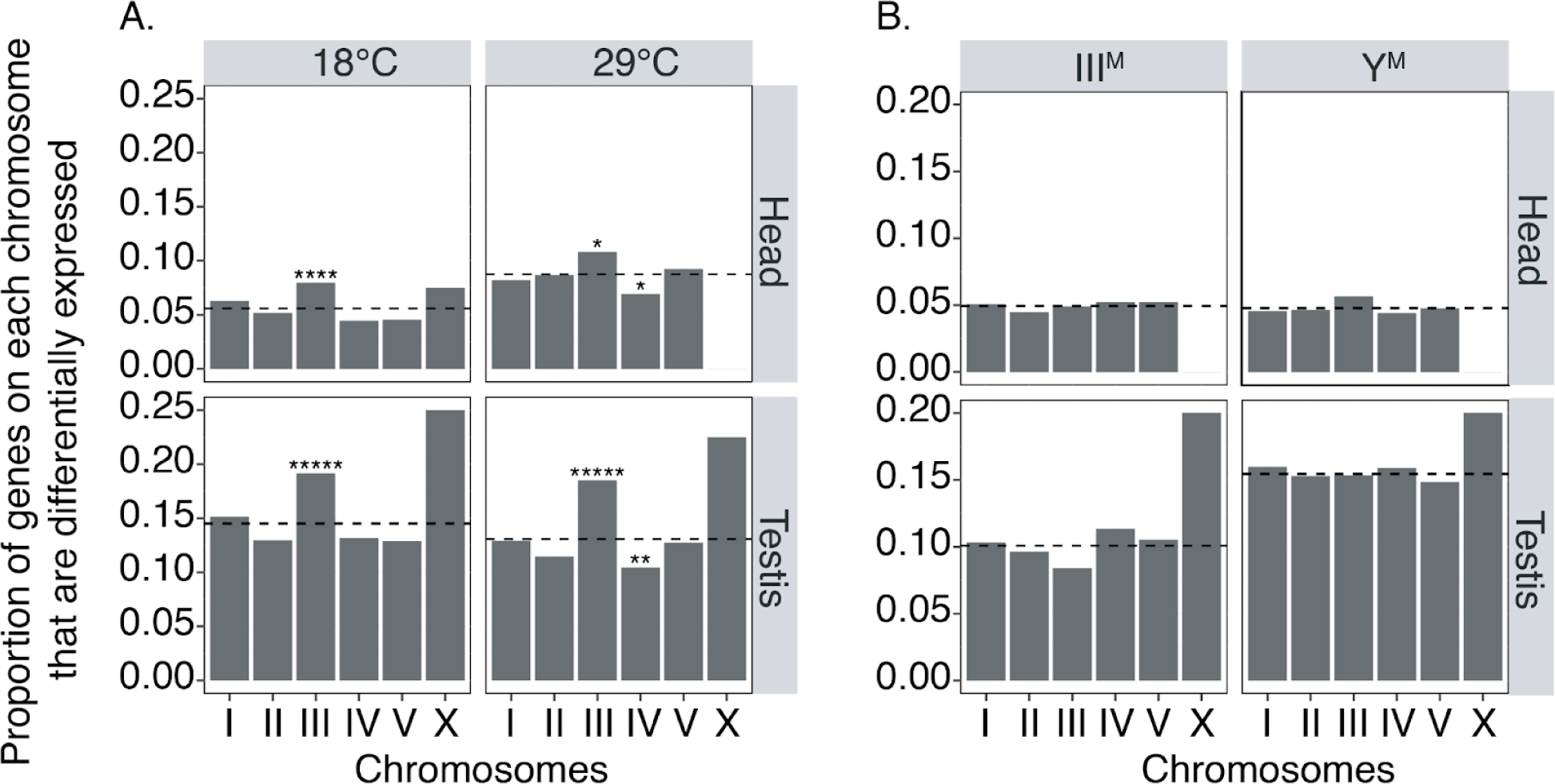
Genes that are differentially expressed between house fly genotypes are significantly enriched on the third chromosome. A) The proportion of house fly genes on each chromosome that are differentially expressed (DE) between Y^M^ and III^M^ males is plotted for heads (top) and testes (bottom) of flies raised at 18°C (left) or 29°C (right). B) The proportion of house fly genes on each chromosome that are DE between temperatures are plotted for heads (top row) and testes (bottom row) for III^M^ (left) and Y^M^ (right) males separately. Each bar represents the proportion of DE genes on a chromosome (# DE genes / # genes on the chromosome), and dashed lines show the the proportion of DE genes across the genome (# DE genes / # genes assigned to any chromosome). Asterisks indicate *P* values obtained from Fisher’s exact test comparing the number of DE genes on a chromosome, the number of non-DE genes on a chromosome, and the number of DE and non-DE genes across all other chromosomes, after Bonferroni correction (\**P* < 0.05, \*\**P* < 0.005, \*\*\**P* < 0.0005, \*\*\*\**P* < 0.00005, \*\*\*\*\**P* < 0.000005).

When comparing between temperatures within each genotype, we found 739 genes significantly differentially expressed between heads of Y^M^ flies raised at the two different temperatures (Supplementary Table 4, Supplementary Figure S7). Similarly, 744 genes are differentially expressed between the heads of III^M^ flies raised at different temperatures (Supplementary Table 4, Supplementary Figure S7). When we excluded older samples, we found 828 genes differentially expressed between heads of III^M^ males raised at different temperatures and 1280 genes differentially expressed between the heads of Y^M^ males (Supplementary Table 5, Supplementary Figure S7). Once again, the increase in differentially expressed genes when the older samples are excluded can be explained by greater power to detect differential expression when the outlier III^M^ males are removed. This also increases power to detect differences within Y^M^ males because we analyze the data with a statistical model that includes all genotypes, temperatures, and their interactions. In testis, there are 2402 genes in Y^M^ flies and 1649 genes in III^M^ flies that are differentially expressed between 18°C and 29°C (Supplementary Table 6, Supplementary Figure S7).

There is no significant chromosomal enrichment of genes that are differentially expressed between temperatures in either head or testis when we include all samples (Figure 2B), consistent with these comparisons being between flies with the same genotype. However, we found a modest enrichment of third chromosome genes that are significantly differentially expressed between temperatures in young Y^M^ heads, i.e., excluding the two older samples (Supplementary Figure S8). This is surprising because all Y^M^ males should have the same third chromosome genotype, and we do not have an explanation for this pattern. As above, the small number of genes on the house fly X chromosome greatly reduces our power to detect significant differences between observed and expected proportions (Meisel & Scott 2018).

### G×T interactions affect the expression of a small subset of genes

We next identified individual genes that are differentially expressed between Y^M^ and III^M^ males depending on temperature by testing for significant interactions between genotype and temperature on gene expression levels. We found 50 genes in head and 247 genes in testis whose expression significantly differs in response to the G×T interaction when we include all samples (Supplementary Tables 4 and 6, Supplementary Figure S7). We found 108 genes differentially expressed in heads in response to the GxT interaction when the two older samples were excluded (Supplementary table 5, Supplementary Figure S7). Of the genes for which the G×T interaction significantly affects expression in head, there are 26 genes that are shared by the analysis of all heads and when the two older samples are excluded (Supplementary Figure S9). We did not find an enrichment of genes with significant G×T interactions on any chromosome in all male heads, younger male heads, or testes (Supplementary Figure S10).

There are 10 genes that are affected by G×T interactions in both head and testis (Supplementary Figure S9). We would expect <1 gene to be affected by G×T interactions in both head and testis if the G×T effects are independent across tissues. The ten genes we observed are significantly greater than this expectation (*z* = 10.12, *P* < 2.2e-16, in a test of proportions), suggesting G×T effects on expression are not independent across tissues. We found 9 genes that are affected by G×T interactions in both testis and young male heads (Supplementary Figure S9), which is significantly greater than the expectation of <2 genes (*z* = 5.42, *P* = 5.88e-08, in a test of proportions). We also found an excess of genes that are differentially expressed in both head and testis in all pairwise comparisons between genotypes and temperatures (Supplementary Figure S9). A similar non-independence of expression differences across tissues was previously observed between Y^M^ and III^M^ males (Meisel *et al*. 2015).

We characterized the functional annotations of genes that are differentially expressed as a result of G×T interactions. We did not find any GO terms associated with genes significantly differentially expressed as a result of GxT interactions in either testis or head, regardless of whether we include all head samples or exclude the two older samples. However, individual genes are suggestive of biological functions that could be affected by GxT interactions on expression. In head, the genes that were differentially expressed because of G×T interactions include an apolipoprotein-D gene (*LOC101893129*). This gene is homologous to *D. melanogaster NLaz*, which is involved in stress response (Hull-Thompson *et al*. 2009), and it is upregulated in III^M^ males at 29°C (Figure 3A). Two genes encoding immune effectors (*LOC105261620,* which encodes a Defensin; and *LOC101895951*, which encodes a Lysozyme and is homologous to *D. melanogaster LysP*) were also upregulated in III^M^ at 29°C (Figure 3A). Three DNA repair genes (*LOC101889156*, homologous to *D. melanogaster Gen*, encoding XPG-like endonuclease; *LOC101899772*, homologous to *maternal haploid*, *mh*, which encodes a protease; and *LOC101899952*, homologous to *Stromalin*, *SA*) are upregulated in Y^M^ at 18°C (Figure 3A). Lastly, an odorant binding protein-coding gene (*LOC105261913*, homologous to *D. melanogaster Obp56h*) was upregulated in Y^M^ males at 29°C (Figure 3A).

**Figure 3.**
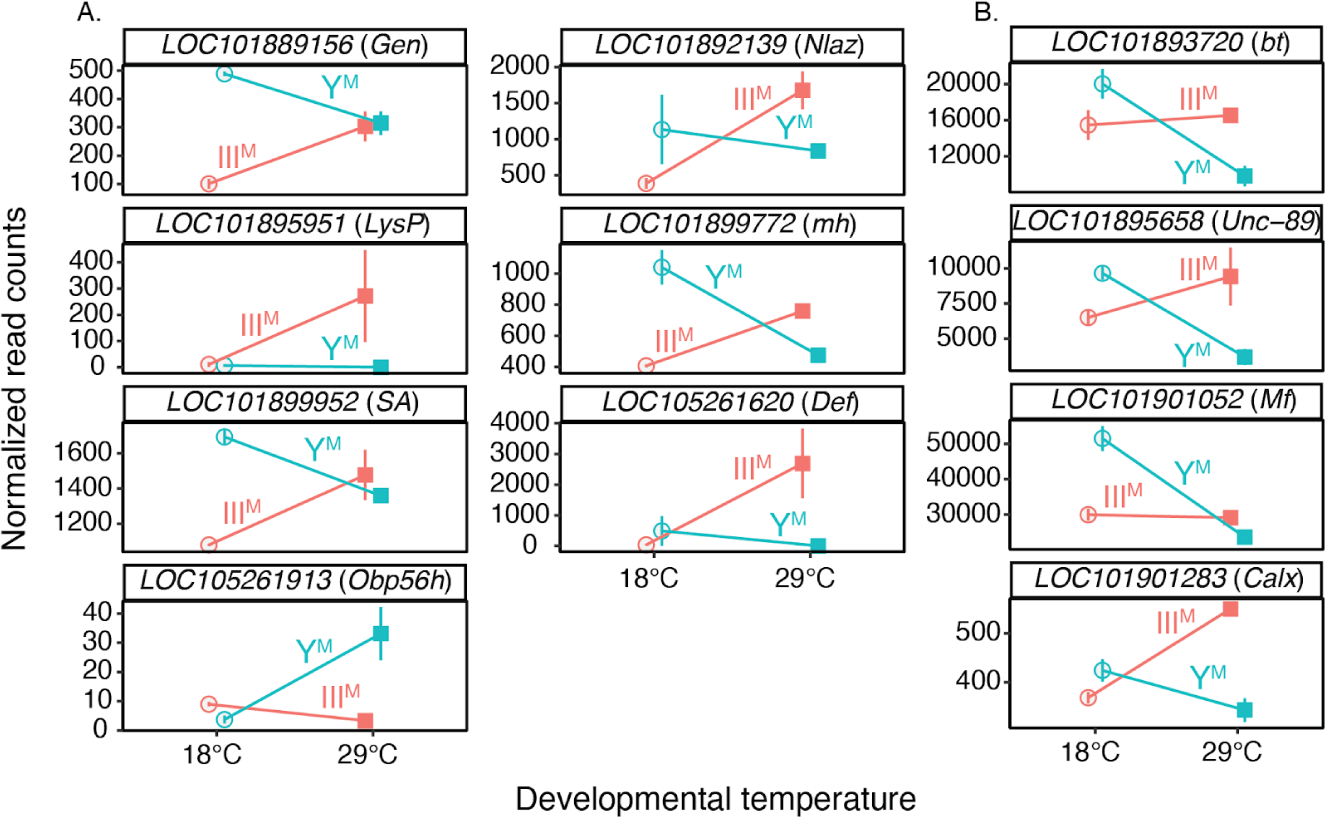
G×T interactions affect gene expression in house fly head. Graphs show normalized read counts (obtained using DESeq2) for genes significantly differentially expressed because of G×T interactions in A) all male head samples or B) young male heads only. Genes are identified based on their house fly gene ID followed by their *Drosophila melanogaster* homologs in parenthesis. Error bars represent standard errors of the mean.

We also identified genes whose expression depends on the G×T interaction when we exclude the two older head samples. *LOC101895951* (*LysP*), *LOC105261913* (*Obp56h*), *LOC101889156* (*Gen*), *LOC101899772* (*mh*), and *LOC101899952* (*SA*) were also significantly differentially expressed in younger heads in the same direction as when we analyze all male head samples (Supplemental Figure 15). A similar pattern was observed for *Nlaz* expression when we only include young heads, although the G×T effect is not significant (Supplementary Figure 15). Other genes only have significant G×T effects in young male heads, including three genes related to muscle performance (*LOC101893720*, homologous to *D. melanogaster bent, bt*; *LOC101895658*, homologous to *Unc-89*; and *LOC101901052*, homologous to *Myofilin*, *Mf*), which are all upregulated in Y^M^ at 18°C (Figure 3B). One gene involved in endoplasmic reticulum (ER) stress response (*LOC101901283*, homologous to *Calx*) is upregulated in III^M^ males at 29°C (Figure 3B).

**Table 1:**
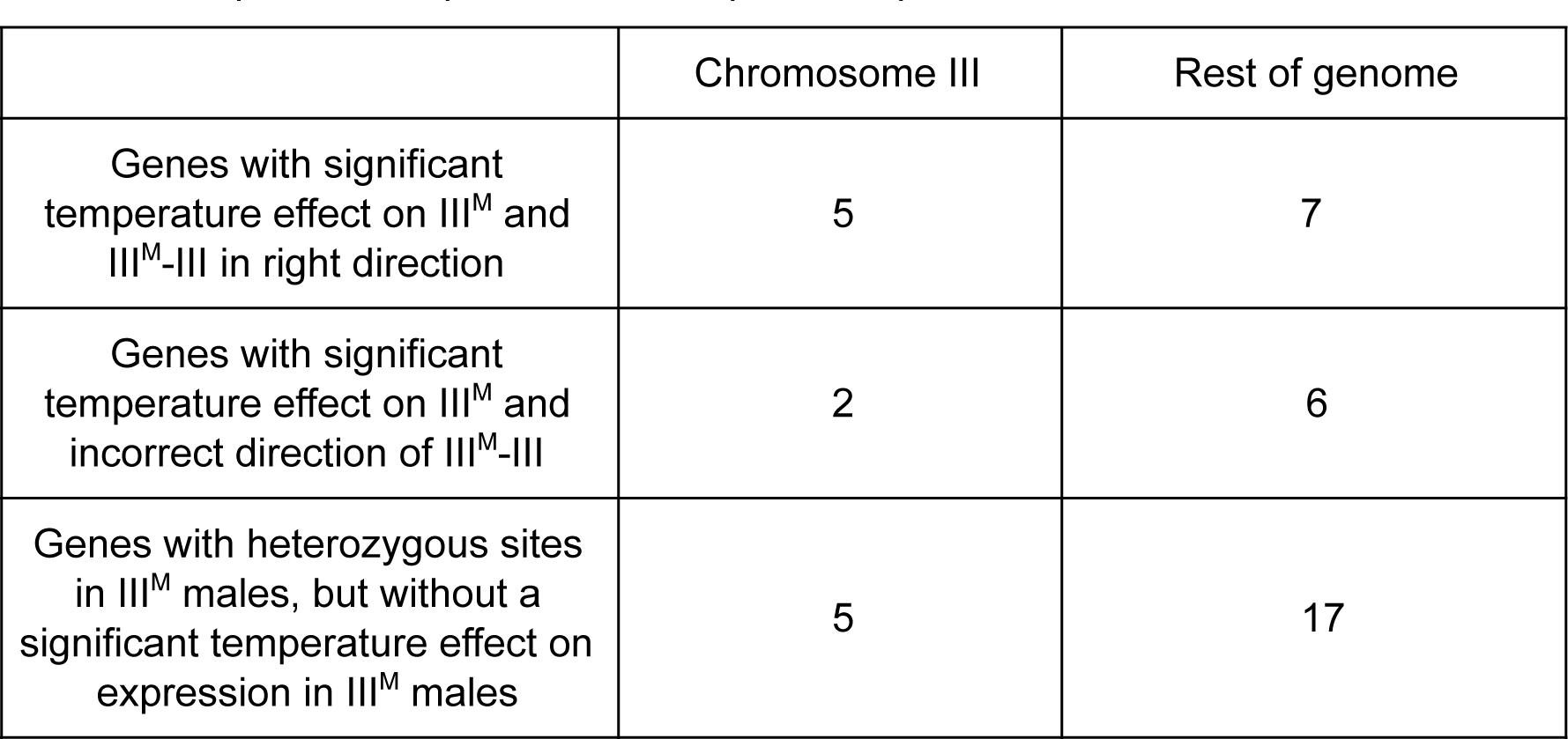
Temperature-dependent allele-specific expression in testis.

|  | Chromosome III | Rest of genome |
| --- | --- | --- |
| Genes with significant temperature effect on $III^M$ and $III^M$-$III$ in right direction | 5 | 7 |
| Genes with significant temperature effect on $III^M$ and incorrect direction of $III^M$-$III$ | 2 | 6 |
| Genes with heterozygous sites in $III^M$ males, but without a significant temperature effect on expression in $III^M$ males | 5 | 17 |

In testis, genes with significant G×T effects on expression include those coding for proteins related to reproductive functions: the protamine *ProtB* homolog *LOC101887804*; the *asunder* (*asun*) homolog *LOC101899763*; the *sarah* (*sra*) homolog *LOC101894442*, and the *Farnesyl pyrophosphate synthase* (*Fpps*) homolog *LOC101896699* (Figure 4). Differential expression of reproduction related genes is expected because testis is the largest male reproductive organ in house fly. Other notable genes that are differentially expressed in testis because of G×T interactions include three metabolic genes (*LOC109613297*, which encodes a hexokinase and is homologous to *D. melanogster Hex-t2*; *LOC101901027,* which encodes fructose-1,6-bisphosphatase and is homologous to *D. melanogaster fbp*; and *LOC101901154*, which encodes an aldehyde oxidase, homologous to *AOX3*), all of which are upregulated in Y^M^ males at 18°C (Figure 4). We also identify one adult lifespan related gene (*LOC101897626*, the homolog of *D. melanogaster pointed*, *pnt*) that is downregulated in III^M^ males at 29°C, and another lifespan related gene (*LOC101897352*, which encodes cystathionine β-synthase, *Cbs*) that is upregulated in Y^M^ males at 18°C (Figure 4). Lastly, two immunity-related genes are differentially expressed in testis. One of the immune genes (*LOC101887442*, which encodes a Gram-negative bacteria-binding protein and is homologous to *GNPB3*) is upregulated in Y^M^ males at 18°C, and the other (*LOC101895929*, which is homologous to *D. melanogaster Phenoloxidase 1*, *PPO1*) is upregulated in III^M^ males at 18°C (Figure 4).

**Figure 4.**
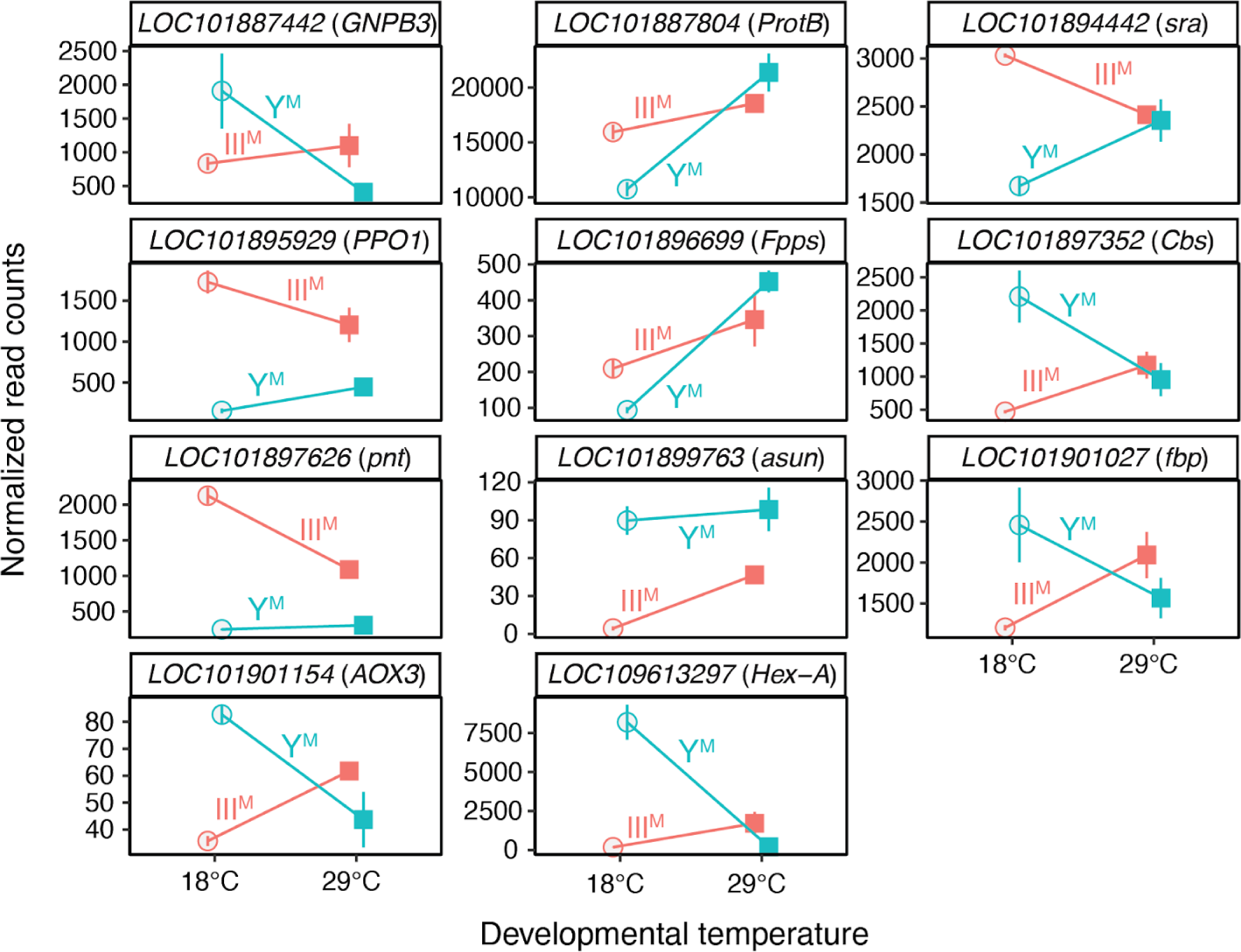
G×T interactions affect gene expression in house fly testis. Graphs show normalized read counts (obtained using DESeq2) for genes significantly differentially expressed in testis because of G×T interactions. Genes are identified based on their house fly gene ID followed by their *Drosophila melanogaster* homologs in parenthesis. Error bars represent standard errors of the mean.

### G×T interactions affecting expression of genes in the sex determination pathway

We did not find evidence that the sex determining gene *Md-tra* is differentially expressed according to a G×T interaction in either all male heads (Supplementary Table 4), young male heads (Supplementary Table 5), or testes (Supplementary Table 6). This suggests that temperature-dependent misregulation of the sex determination pathway is not responsible for fitness differences of Y^M^ and III^M^ males across the cline. In comparison, we found evidence of effects of G×T interactions on the expression of most *Md-tra* exons in both head (including or excluding older samples) and testis (Supplementary Figures S11 and S12). However, if a G×T interaction affecting the mis-splicing of *Md-tra* were responsible for the latitudinal distribution of Y^M^ and III^M^, we would expect more female-determining isoforms produced (i.e., misexpressed) in Y^M^ males raised at a high temperature, or a higher expression level of female-determining isoforms in III^M^ males raised at a low temperature. In contrast to that expectation, the G×T interactions are not in the directions consistent with misexpression at discordant temperatures (Supplementary Figure S11 and S12). An analysis of *Md-tra* splicing with qPCR was not possible because we could not design primers that specifically amplified isoforms for quantitative assessment.

We further tested if G×T interactions affect the expression and splicing of two direct downstream targets of *Md-tra* in the sex determination pathway, *Md-dsx* and *Md-fru*. Our RNA-seq data show that there is no effect of G×T interactions in the expression of *Md-dsx* or *Md-fru* in all male heads (Supplementary Table 4), young male heads (Supplementary Table 5), or testes (Supplementary Table 6). We also found no evidence of G×T interactions affecting the expression of individual *Md-dsx* exons (Supplementary Figure S13). We did not test for G×T effects on the expression of *Md-fru* exons because exons that differentiate the male and female isoforms have not been annotated in the reference genome (Meier *et al*. 2013; Scott *et al*. 2014).

We also used qRT-PCR to examine the expression of the house fly male-determining gene, *Mdmd*, in two III^M^ strains and two Y^M^ strains raised at 18°C and 27°C (Supplementary Figure S14). One Y^M^ strain and one III^M^ strain originated from North America, and the other Y^M^ strain and III^M^ strain came from Europe. If temperature-dependent differential expression of *Mdmd* were responsible for the clinal distribution of Y^M^ and III^M^ (with higher expression conferring a fitness advantage), we would expect higher *Mdmd* expression in Y^M^ (III^M^) males at lower (higher) temperatures. There is a significant G×T interaction affecting the expression of *Mdmd* in the European Y^M^ and III^M^ strains, with higher *Mdmd* expression in III^M^ males at lower temperatures (Supplementary Figure S14). This is the opposite pattern from what would be expected if the hypothesized G×T effects on *Mdmd* expression were responsible for maintaining the cline. A similar trend is observed in the North American strains, although the interaction term is not significant. We observe these similar patterns in both population samples even though they were assayed with two different types of tissue (abdomen in the North American strains, and whole fly in the European strains), demonstrating that these results are robust to the tissues we sampled. We also did not find a significant G×T interaction affecting expression of *Md-ncm* (the ancestral paralog of *Mdmd*), which is not part of the sex determination pathway (Supplementary Figure S14). Therefore, there is no evidence that *Mdmd* expression is increased at the hypothesized favored temperatures for Y^M^ and III^M^ males.

### G×T interactions on gene expression are not driven by cis-regulatory divergence

We next tested if divergence of *cis*-regulatory sequences between the III^M^ and standard third chromosome is responsible for temperature-dependent expression differences between III^M^ and Y^M^ males. III^M^ males are heterozygous (III^M^/III) whereas Y^M^ males are homozygous (III/III) for a standard third chromosome. If *cis*-regulatory alleles on the third chromosome are responsible for differential expression of third chromosome genes between III^M^ and Y^M^ males, the III^M^ and III alleles of those genes should also be differentially expressed in III^M^ males. For example, if a gene is more highly expressed in III^M^ males than Y^M^ males, the III^M^ allele of the gene should be more highly expressed than the III allele in III^M^ males. The opposite would be true if Y^M^ males have higher expression than III^M^ males. We used this logic to test if G×T interactions on gene expression are the result of *cis-*regulatory divergence of third chromosome genes between the III^M^ and III chromosomes. To do so, we asked if genes on the third chromosome that are significantly differentially expressed in head or testis because of G×T interactions have concordant differences in expression between the III^M^ and III allele in III^M^ males.

To test for differences in allelic expression, we first identified 12 genes on the third chromosome with a significant G×T interaction affecting testis expression, at least one heterozygous SNP in III^M^ males, and homozygous at those SNP sites in Y^M^ males (Figure 5). We required the variants to be heterozygous in III^M^ males and homozygous in Y^M^ males because we are interested in expression differences between the III^M^ and III allele in III^M^ males. We assumed that the allele in common between III^M^ and Y^M^ males is found on the standard third chromosome, and the allele unique to III^M^ males is on the III^M^ chromosome. This assumption is reasonable because the Y^M^ and III^M^ flies that we used for our RNA-seq share the same genetic background, and therefore should have the same standard third chromosome. We quantified the expression of the two alleles (III^M^ and III) based on allele-specific RNA-seq read coverage. We asked if the difference in expression of III^M^ alleles in each gene is consistent with the difference in overall expression of these genes between 18°C and 29°C within III^M^ males. For example, if III^M^ males have higher expression at 29°C, we expect the difference between the III^M^ and III alleles to be greater at 29°C than 18°C.

**Figure 5:**
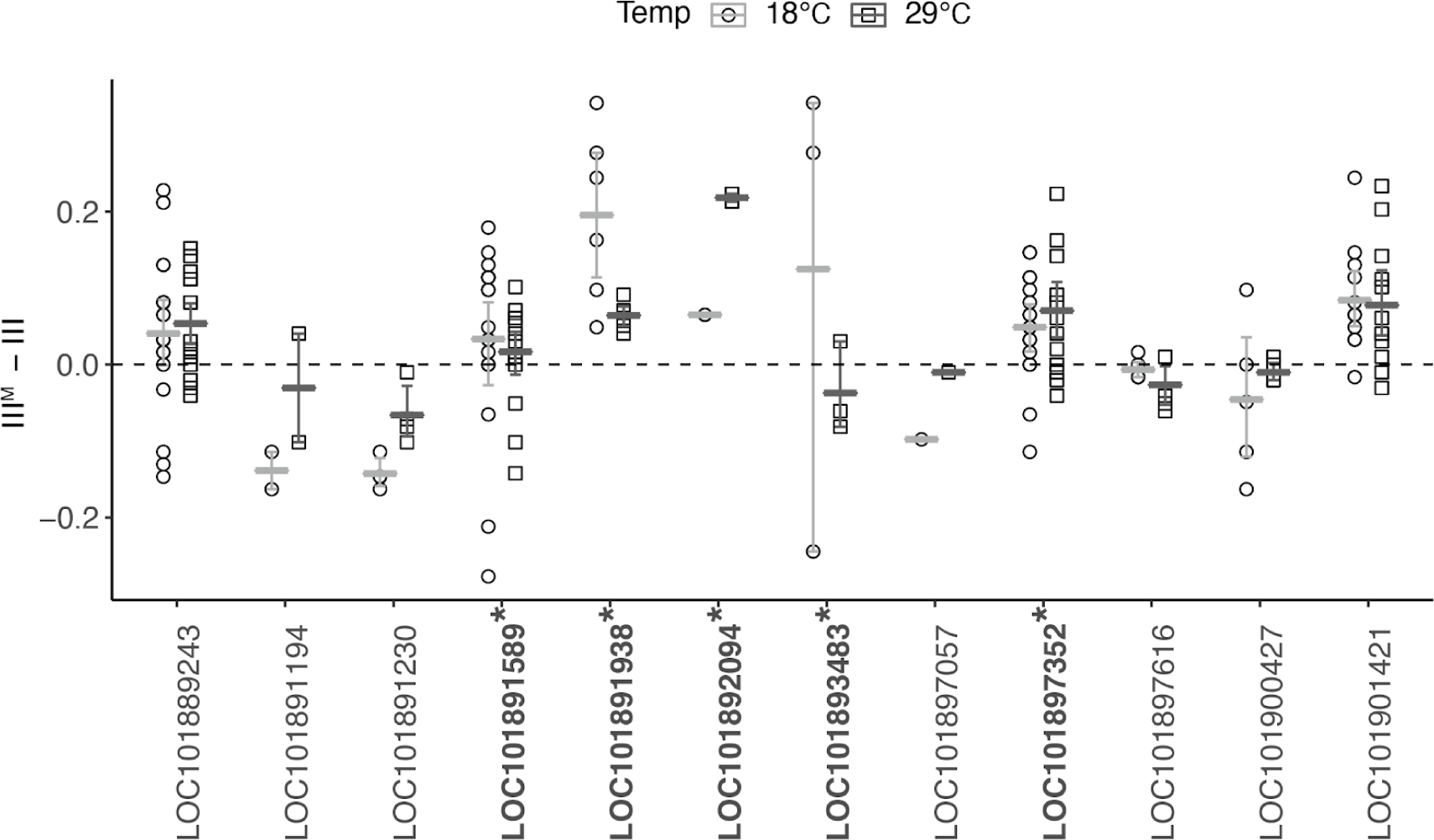
G×T interactions affect allele-specific expression in house fly testes. Differences in sequencing coverage in III^M^ house fly males between III^M^ and III alleles at either 18°C (circles) or 29°C (squares) are shown for 12 house fly genes where there is a G×T effect on testis expression between Y^M^ and III^M^ males. Each circle or square represents the difference in normalized mapped reads in testis between the III^M^ and III alleles at a single variable site (SNP) within a gene. Circles show expression differences between alleles at 18°C, and squares show expression differences between alleles at 29°C. The small horizontal lines indicate the mean difference in coverage between alleles across all sites in each gene at each temperature. Error bars represent the standard error across all variable sites within a gene at each temperature. The gene names with asterisks along the x-axis are differentially expressed between the III^M^ and III alleles in the same direction as the differential expression in III^M^ males between 18°C and 29°C (Table 1).

We first compared the expression of III^M^ and III alleles in testis. Of the 12 genes with significant G×T effects and the requisite SNPs to test for allele-specific expression, seven have a significant effect of temperature on testis gene expression within III^M^ males (Figure 5; Table 1). Of those seven genes, five have a pattern of allelic expression consistent with the differential expression between 18°C and 29°C within III^M^ males: *LOC101892094* (homologous to *D*. *melanogaster Pdfr*, which is responsible for regulating circadian behaviors), *LOC101891589* (homologous to *D*. *melanogaster CG42450*, which is predicted to be involved in G protein-coupled receptor signaling), *LOC101893483* (encoding a GATA zinc finger domain-containing protein), *LOC101891938* (homologous to *D. melanogaster mmd*, which is predicted to encode a membrane protein involved in ectodomain proteolysis), and *LOC101897352* (the cystathionine β-synthase gene associated with lifespan, mentioned earlier). The two genes with allelic expression that is inconsistent with temperature-dependent expression in III^M^ males are *LOC101882943* (homologous to *D. melanogaster Nepl5*) and *LOC101900427* (homologous to *D. melanogaster fne*). The remaining five genes do not differ in testis expression between III^M^ males raised at 18°C and 29°C (Table 1).

To determine a null expectation for the proportion of genes with allelic expression consistent with the differential expression between 18°C and 29°C, we tested for concordance between allele-specific expression and temperature-dependent expression differences for genes on other chromosomes. We do not expect concordance for genes on other chromosomes because the inbred Y^M^ and III^M^ males used in our RNA-seq experiment share a common genetic background. We identified 30 genes on other chromosomes with heterozygous sites whose testis expression depends on the G×T interaction (Table 1). Of those 30 genes, 13 are differentially expressed between III^M^ males raised at 18°C and 29°C. We find that 7 out of the 13 genes in the rest of the genome have allele-specific expression that is consistent with the 18°C vs 29°C expression differences (Table 1). There is not a significant excess of genes on the third chromosome whose temperature-dependent expression is consistent with changes in allele-specific expression relative to the rest of the genome (*P* = 0.64 in Fisher’s exact test). This suggests that the G×T effects on the expression of genes on the third chromosome is not the result of an excess of *cis*-regulatory differences between the III^M^ and standard third chromosomes.

When we analyzed only the younger male head samples, we found 7 genes on the third chromosome with a significant G×T interaction that also had at least one SNP in III^M^ males. Among them, one gene (*LOC101890343*, homologous to *D. melanogaster mahe*, encoding an ATP-dependent RNA helicase) had a significant effect of temperature on gene expression within III^M^ males. The allele specific expression of this gene is consistent with the temperature effect in III^M^ males, but there are no genes on other chromosomes with the requisite SNPs in our head RNA-seq data to test for a significant excess relative to a null expectation. When analyzing all head samples, we found a single gene on the third chromosome with a significant GxT interaction that also had a SNP in III^M^ males. However, we did not find a significant effect of temperature on expression of this gene within III^M^ males.

We are limited in the analysis we can perform on allele-specific expression of genes on the X vs Y^M^ chromosomes because of small sample sizes. There are only 40 genes assigned to the house fly X or Y^M^ chromosome (Meisel & Scott 2018), none of which have a significant G×T interaction affecting expression in testis (Supplementary Table 3). Only one X or Y^M^ chromosome gene has a significant G×T interaction affecting expression in heads when we analyze all samples (Supplementary Table 2), and it does not have any heterozygous sites. Similarly, none of the three genes on the X chromosome with a significant G×T interaction affecting expression in young male heads has any heterozygous sites.

## Discussion

We tested for G×T interactions that affect gene expression in Y^M^ and III^M^ house fly males. These G×T effects could lead to differences in temperature-dependent phenotypes between house fly genotypes, thereby maintaining polygenic sex determination across latitudinal clines based on temperature-dependent fitness effects of the proto-Y chromosomes. We used RNA-seq to compare gene expression in heads and testes of two nearly isogenic strains that differ only in their proto-Y chromosomes (Y^M^ or III^M^) that we raised at two different temperatures (18°C and 29°C). This 2×2 full factorial design allowed us to compare genome-wide expression between four G×T combinations, which we combined with targeted expression measurements of the male-determining gene (*Mdmd*) using qRT-PCR. We found that G×T interactions lead to differential gene expression in both head and testis, but the expression of genes involved in the sex determination pathway is not meaningfully affected by those G×T interactions. We therefore hypothesize that alleles present on either the III^M^ chromosome or the Y^M^ chromosomes, other than *Mdmd*, may be targets of selection.

### No evidence that G×T interactions affect the sex determination pathway in a way that explains the maintenance of polygenic sex determination

Our results suggest that G×T interactions affecting the sex determination pathway do not explain the maintenance of polygenic sex determination in house fly. Evolutionary transitions between heritable and temperature-dependent sex determination systems are possible if sex determination pathways are temperature sensitive (Shine *et al*. 2002; Quinn *et al*. 2007; Radder *et al*. 2008; Holleley *et al*. 2015). Sex determination in flies operates by alternative splicing of multiple genes in the pathway (Salz 2011; Bopp *et al*. 2014). Temperature dependent alternative splicing has been reported in *Arabidopsis* (Streitner *et al*. 2013; Steffen & Staiger 2017), *Neurospora* (Colot *et al*. 2005), *Drosophila* (Jakšić & Schlötterer 2016; Martin Anduaga *et al*. 2019), and mammals (Preußner *et al*. 2017). It is therefore possible that temperature-sensitive expression or splicing of sex determination factors can establish a clinal distribution of sex determination genes, such as what is observed in house fly (Schenkel 2021). We did not find evidence for G×T interactions affecting the expression of the male-determining *Mdmd* gene or splicing of *Md-tra* in a way that is consistent with the clinal distribution of Y^M^ and III^M^. In addition, the expression of *Md-dsx* and *Md-fru*, the immediate downstream targets of *Md-tra*, do not depend on G×T interactions.It is possible that temperature affects the expression or splicing of sex determination pathway genes earlier in development than we measured. For example, *Mdmd* expression level might be more critical during early embryogenesis when *Md-tra* needs to be locked into a male or female mode of splicing (Sharma *et al*. 2017). Hediger et al. (2010) have shown that the *Md-tra* auto-regulatory loop can be effectively shut down in embryos by RNA interference, and male development proceeds normally without the need of *Mdmd* expression. Similarly, when *Mdmd* was removed from *Mdmd-*/+ cells at embryonic stages, the resulting clones developed as males despite their female genotype (Hilfiker-Kleiner *et al*. 1993). Thus the adult *Mdmd* and *Md-tra* expression we observed might not reflect the critical early expression levels. Additional work is required to further examine temperature-dependent effects on the expression or splicing of *Mdmd* or *Md-tra* across male genotypes in embryos, larvae, or pupae, rather than in adults.

Even though we did not observe differential expression of *Mdmd* that is consistent with our hypothesis for the clinal distribution of Y^M^ and III^M^ males, we believe that the increased expression of *Mdmd* in III^M^ males that we observe at the lower temperature is intriguing. It is possible that *Mdmd* expression is optimal at an intermediate level between high and low extremes—lower expression of *Mdmd* might be insufficient for *Md-tra* splicing, whereas higher expression of *Mdmd* might be toxic because of its proposed role in antagonizing functions of the generic splicing factor *Md-ncm* (Sharma *et al*. 2017). The increased expression of *Mdmd* in III^M^ males at a lower temperature might thus explain the absence of III^M^ males in northern latitudes. Moreover, Hediger et al. (1998) found male determining regions on both arms of the Y^M^ chromosome that act additively. However, it is not yet resolved whether *Mdmd* is the male determining factor on both of these arms or only one arm (Sharma *et al*. 2017). Additional work is required to determine if there is an additional male determining gene other than *Mdmd* on the Y^M^ chromosome that may have temperature dependent activity.

### Temperature-dependent gene expression is not the result of large-scale cis-regulatory changes on the III^M^ chromosome

Previous RNA-seq experiments (Meisel *et al*. 2015; Son *et al*. 2019), as well as the results presented here (Figure 2A), provide consistent evidence that the third chromosome is enriched for genes that are differentially expressed between Y^M^ and III^M^ males. This is expected as the comparisons are between flies that differ in their third chromosome genotypes, and it suggests there are *cis*-regulatory effects on the expression of genes on the third chromosome. Consistent with this hypothesis, we observe a more pronounced clustering by genotype in our PCA when we consider only chromosome III genes (Supplementary Figure S4). In contrast, we find that genes that are differentially expressed because of temperature are not enriched on the third chromosome in III^M^ males (Figure 2B). This is not because of lack of power to detect the enrichment as we see a modest enrichment of differentially expressed third chromosome genes in young Y^M^ male heads (Supplementary Figure 8). We also found that G×T interactions affecting the expression of genes in male heads or testes are not enriched on the third chromosome either (Supplementary Figure S9).

The lack of an enrichment of genes with temperature-dependent expression in III^M^ males on the third chromosome suggests that temperature-dependent effects of the III^M^ chromosome are not mediated by large-scale *cis-*regulatory changes across the III^M^ chromosome. Consistent with this interpretation, there is not an enrichment of third chromosome genes with temperature-dependent expression differences between the III^M^ and III alleles (Figure 5, Table 1). Moreover, an independent analysis of other RNA-seq data also found that there is not an excess of expression differences between III^M^ and III alleles in a different house fly strain (Son & Meisel 2021). We cannot perform a similar statistical analysis of Y^M^ genes because of the small number of genes on that chromosome.

### Temperature-dependent gene expression and the maintenance of polygenic sex determination in house fly

We hypothesized that targets of selection responsible for the maintenance of polygenic sex determination in house fly could be differentially expressed across proto-Y chromosome genotypes and developmental temperatures. Despite our conclusion that a large number of *cis-*regulatory variants on the III^M^ chromosome cannot explain the effect of the III^M^ chromosome on temperature-dependent phenotypes, we still found evidence for temperature-dependent effects of the III^M^ and Y^M^ chromosomes that could explain their divergent phenotypic effects. First, there is some clustering by G×T combinations in the transcriptome-wide testis gene expression profiles (Figure 1C). Second, we identify substantial temperature-dependent gene expression (Figure 2B) and many genes whose expression depend on G×T interactions (Figures 3 and 4). These temperature-dependent effects on expression could be responsible for phenotypic differences between Y^M^ and III^M^ males, which could in turn provide a substrate upon which selection acts to maintain the Y^M^-III^M^ clines. If wide-spread *cis*-regulatory differences across proto-sex chromosomes are not responsible for these G×T effects (as we hypothesize above), then it is possible that a small number of loci on the proto-Y chromosomes act as temperature-dependent *trans* regulators of gene expression across the entire genome.

Reproductive traits are a promising target of selection that could depend on G×T interactions. There are more genes differentially expressed in testis because of G×T interactions than in head, consistent with previous work that identified more differentially expressed genes in testis than head between Y^M^ and III^M^ males (Meisel *et al*. 2015). Genes associated with reproductive functions (*LOC101887804*, *LOC101899763*, *LOC101894442*, and *LOC101896699*) were amongst the genes whose testis expression depend on G×T effects (Figure 4). It is therefore possible that selection along the Y^M^-III^M^ cline acts on reproductive traits, which is consistent with the idea that the strength of sexual selection can vary across populations (Arnqvist 1992; Payne & Krakauer 1997; Blanckenhorn *et al*. 2006; Connallon 2015; Allen *et al*. 2017). These reproductive traits, or other variants under selection, could have sexually antagonistic fitness effects (i.e., opposing fitness effects in males and females) which may be temperature-sensitive. Sexual antagonism is one of the few selection pressures capable of maintaining polygenic sex determination (Rice 1986; van Doorn & Kirkpatrick 2007). Population genetic modeling also predicts that sexually antagonistic effects of Y^M^ and III^M^ can maintain polygenic sex determination within house fly populations (Meisel *et al*. 2016; Meisel 2021), possibly in conjunction with epistatic interactions between either Y^M^ or III^M^ and autosomal loci not linked to either *Mdmd* locus (Schenkel 2021). It is worth pursuing if sexual antagonism can maintain polygenic sex determination by acting on temperature-dependent gene expression differences between Y^M^ and III^M^ males.

Energy metabolism is a potential phenotype upon which selection acts to affect reproductive functions. We previously found divergence between III^M^ and standard third chromosome sequences surrounding genes encoding mitochondrial proteins (Son & Meisel 2021). Here, we report G×T interactions affecting the testis expression of three genes with metabolic functions (*LOC101901027*, *LOC101901154*, and *LOC109613297*). All three genes are upregulated in Y^M^ males at 18°C, and, to a lesser extent, upregulated in III^M^ males at 29°C (Figure 4). None of the *D. melanogaster* homologs of these genes are differentially expressed between flies raised at high (21.5°C) or low (6°C) temperatures (MacMillan *et al*. 2016), nor are they differentially expressed between *D. melanogaster* that are evolved in hot or cold laboratory environments (Hsu *et al*. 2020). However, one of the metabolic genes (*LOC101901154*), encoding an aldehyde oxidase, has a *D. melanogaster* homolog (*AOX4*) that is expressed higher at 21°C than 29°C (Zhao *et al*. 2015), consistent with the higher expression of the house fly gene in Y^M^ males at lower temperatures. We are cautious to interpret further because there are four tandemly arrayed *AOX* genes in the *D. melanogaster* genome and at least 3 corresponding genes in house fly; it is therefore not possible to assign orthology across this family.

One of the other metabolic genes (*LOC101901027*) has a homolog (*fbp*) that is expressed higher in *D. melanogaster* raised at 29°C than those raised at 21°C, regardless of whether the flies come from Maine (USA) or Panama (Zhao *et al*. 2015). This is consistent with the higher expression of this gene at 29°C in III^M^ testes, but opposite from the lower expression at 29°C in Y^M^ testes (Figure 4). It is possible that the Y^M^ chromosome confers a fitness advantage via increased production of fructose-1,6-bisphosphatase in testes at lower temperatures. Consistent with this hypothesis, fructose-1,6-bisphosphatase is necessary for cold-stress in mice (Park *et al*. 2020) and associated with cold hardiness in plants and insects (Storey & Storey 2012; Cai *et al*. 2018). There is also evidence that *D. melanogaster fbp* is differentially *trans-*regulated across genotypes and temperatures (Chen *et al*. 2015). In house fly, this gene is not found on either the Y^M^ or III^M^ chromosome, which would require it to be differentially regulated in *trans*, consistent with what is observed in *D. melanogaster*. Together with the other differentially expressed metabolic genes, our results suggest that energy metabolism related to spermatogenesis or sperm function may be a target of selection driving the evolution of the III^M^ and Y^M^ chromosomes.

Muscle performance might also be under differential selection across the Y^M^-III^M^ cline. We identified three muscle component related genes (*LOC101893720*, *LOC101895658*, and *LOC101901052*) upregulated in Y^M^ male heads at 18°C (Figure 3B). One of these genes (*LOC101893720*) is homologous to *D. melanogaster bt*. Knockdown of *bt* decreases sarcomere length and reduces climbing ability in *D. melanogaster* (Perkins & Tanentzapf 2014). Another muscle-related gene (*LOC101895658*) is homologous to *D. melanogaster Unc-89*, which encodes an obscurin protein. Reduced expression of *Unc-89* using P-element insertion results in flightless adults in *D. melanogaster* (Katzemich *et al*. 2012). Upregulation of these genes in Y^M^ males at lower temperatures might improve muscle performance.

We also find evidence that selection may have acted in response to thermal stress across environments along the Y^M^-III^M^ cline. A gene (*LOC101893129*) homologous to *D. melanogaster Nlaz*, which encodes an extracellular lipid binding protein (similar to apolipoprotein D and Retinol Binding Protein 4), is upregulated in heads of III^M^ males at the high temperature (Figure 3A). *Nlaz* is regulated by the JNK signalling pathway to confer stress and starvation tolerance, and it reduces oxidative stress by maintaining metabolic homeostasis (Hull-Thompson *et al*. 2009). *NLaz* mutants in *D. melanogaster* have reduced stress resistance and shorter lifespans, while over-expressing *NLaz* increases stress tolerance and extends lifespan. *Nlaz* is also upregulated at extreme low temperature in *D. melanogaster* (Chen *et al*. 2015; MacMillan *et al*. 2016). Upregulation of this gene may therefore help III^M^ males tolerate thermal stress at high temperatures. Our results demonstrate the utility of simultaneously studying the effects of both genotypic and temperature variation to determine how thermal stress affects gene expression (Rivera *et al*. 2021).

There is also evidence that improved response to thermal stress may act to increase lifespan in Y^M^ and III^M^ males at temperatures concordant with their clinal distribution. For example, *LOC101897352* encodes cystathionine β-synthase and is homologous to *D. melanogaster Cbs.* In *D. melanogaster*, *Cbs* is involved in ER stress response (Chow *et al*. 2013) and is a positive regulator of lifespan (Kabil *et al*. 2011)*. LOC101897352* is upregulated in Y^M^ male testes at 18°C (Figure 4), consistent with longer lifespan for Y^M^ males at lower temperatures. *LOC101897352* (the *Cbs* homolog) is also one of the genes with a consistent direction of allele-specific expression and expression difference between III^M^ males at 18°C and 29°C (Figure 5), providing evidence that a *cis*-regulatory allele on the III^M^ chromosome drives temperature-dependent expression of a gene with a potential phenotypic effect. Future work should aim to identify *cis*-regulatory regions underlying the temperature-dependent expression differences between the III^M^ and III alleles in *LOC101897352* (the *Cbs* homolog) and other such genes on the third chromosome (Figure 5). Searching for such regulatory sequences in house fly is currently impeded by the lack of a chromosome-scale genome assembly and comprehensive gene annotations (Scott *et al*. 2014; Meisel & Scott 2018).

Two other genes are differentially expressed in a way that is suggestive of temperature-dependent lifespan differences between Y^M^ and III^M^ males. *LOC101895929* is homologous to *D. melanogaster pnt*. Knockdown of *pnt* extends lifespan in *D. melanogaster* (Dobson *et al*. 2019). Interestingly, we see downregulation of this gene in III^M^ male testes 29°C (Figure 4), consistent with longer lifespan for III^M^ males at a higher temperature. Lastly, *LOC101901283* is homologous to *D. melanogaster Calx*, which is associated with response to ER stress (Chow *et al*. 2013)*. Calx* mutation reduces *D. melanogaster* lifespan (Mok *et al*. 2020). *LOC101901283* is upregulated in III^M^ male heads at 29°C (Figure 3B), suggesting a longer lifespan for III^M^ males at a higher temperature. All three lifespan-related genes (*LOC101897352*, *LOC101895929*, and *LOC101901283*) therefore have expression profiles consistent with longer lifespan of Y^M^ males at lower temperatures or III^M^ males at higher temperatures, suggesting that temperature-dependent senescence might be a phenotype under differential selection between III^M^ and Y^M^ males. It remains to be tested if these male genotypes have different lifespans across temperatures.

## Conclusion

The clinal distribution of the house fly proto-Y chromosomes in natural populations hints at a possible G×T interaction involved in maintaining polygenic sex determination (Hiroyoshi 1964; Mcdonald *et al*. 1975; Denholm *et al*. 1986; Hamm *et al*. 2005; Feldmeyer *et al*. 2008; Kozielska *et al*. 2008). We did not find evidence that temperature-dependent expression or splicing of genes in the sex determination pathway explain the maintenance of polygenic sex determination in house fly. However, such effects may act earlier in development than we assayed. In contrast, G×T interactions affect gene expression in both somatic and reproductive tissues across the entire genome. Our results therefore suggest that alleles on the proto-Y chromosomes other than the male-determining *Mdmd* gene are targets of selection responsible for maintaining the proto-Y chromosome clines in house fly.

There is no enrichment of G×T effects on the expression of genes on the proto-Y chromosomes, suggesting that temperature-dependent expression differences between Y^M^ and III^M^ males (and thereby phenotypic and fitness effects of the proto-Y chromosomes) are not driven by a large-number of *cis*-regulatory changes on the III^M^ chromosome. Instead, if temperature-dependent gene expression is responsible for temperature-dependent phenotypic effects of the III^M^ and Y^M^ proto-Y chromosomes, those effects are the result of a small number of alleles on the III^M^ (and possibly Y^M^) chromosome. One such sex-linked gene, encoding cystathionine β-synthase, is differentially expressed across genotypes and temperatures in a way that is consistent with divergence in *cis-*regulatory sequences between the III^M^ and III chromosomes. However, most of the differentially expressed genes between Y^M^ and III^M^ males are not sex-linked and therefore likely the result of *trans* G×T effects on gene expression across the entire genome. This is consistent with our previous work that identified very few differentially expressed genes as a result of differences in proto-Y chromosome genotypes (Son *et al*. 2019). Autosomal genes whose expression depends on proto-Y genotype and temperature include those encoding metabolic proteins in testis, proteins involved in stress response in head, or with effects on aging. This suggests temperature-dependent sperm function or thermal stress tolerance may be targets of selection maintaining the Y^M^-III^M^ cline via *trans*-effects of the proto-Y chromosomes on expression of genes affecting these phenotypes across the genome.

## Acknowledgements

This work was supported by the National Science Foundation (OISE-1444220 and DEB-1845686 to RPM), Mindlin Foundation (MF16-US04 to JJ), and the University of Houston (start up funds to RPM and Provost’s Undergraduate Research Scholarship to JJ). We thank Dalia Aldin and Ashley Lee for assistance establishing the Y^M^ and III^M^ strains with a common genetic background.

## Data Accessibility

RNA-seq data were collected for this manuscript and have been deposited in the NCBI Gene Expression Omnibus under accession GSE136188 (BioProject PRJNA561541, SRA accession SRP219410).

## Supplementary Figures

**Supplementary Figure S1.**
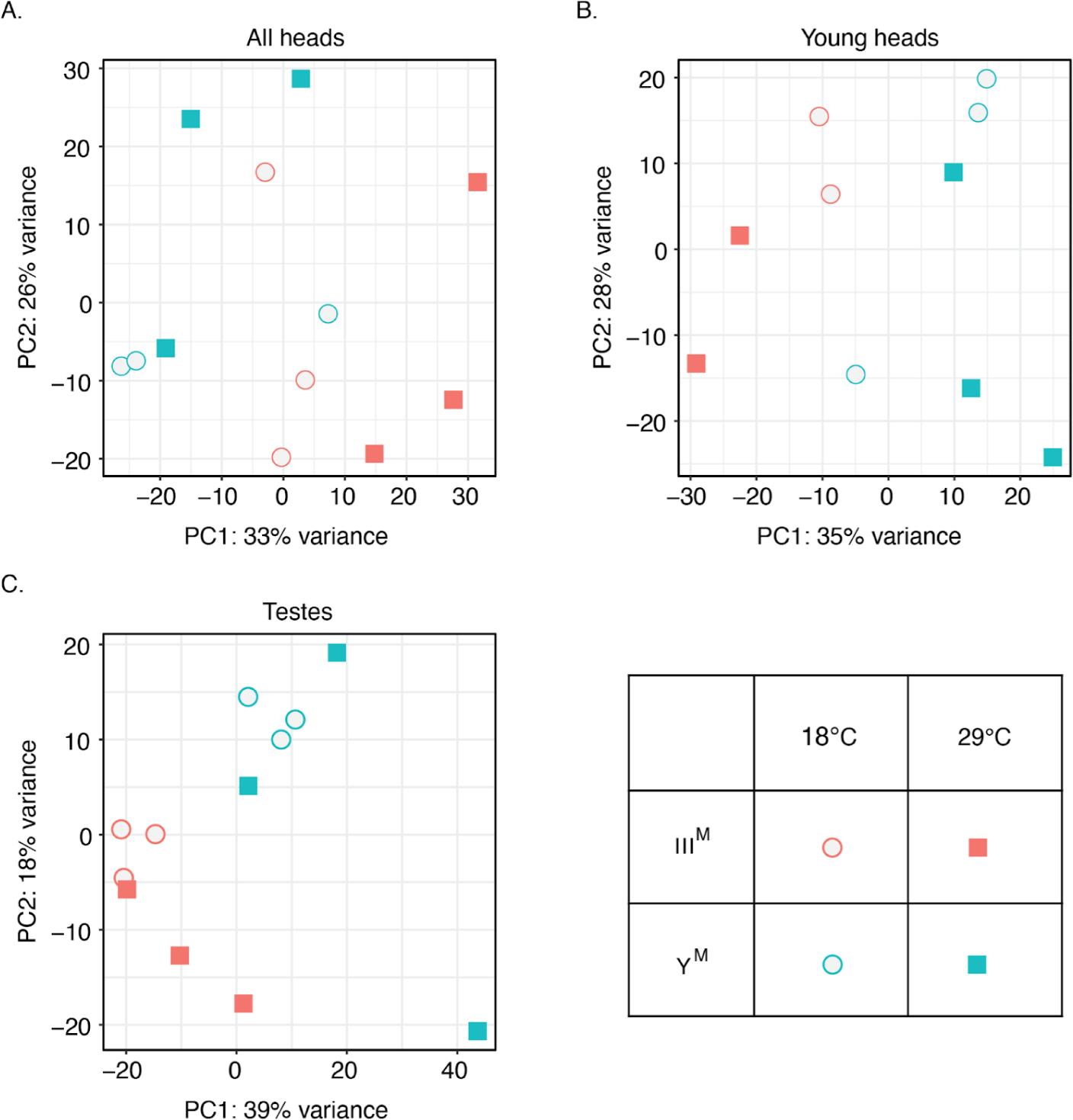
PCA of the 500 most variable genes in A) all male heads, B) young male heads and C) testes.

**Supplementary Figure S2.**
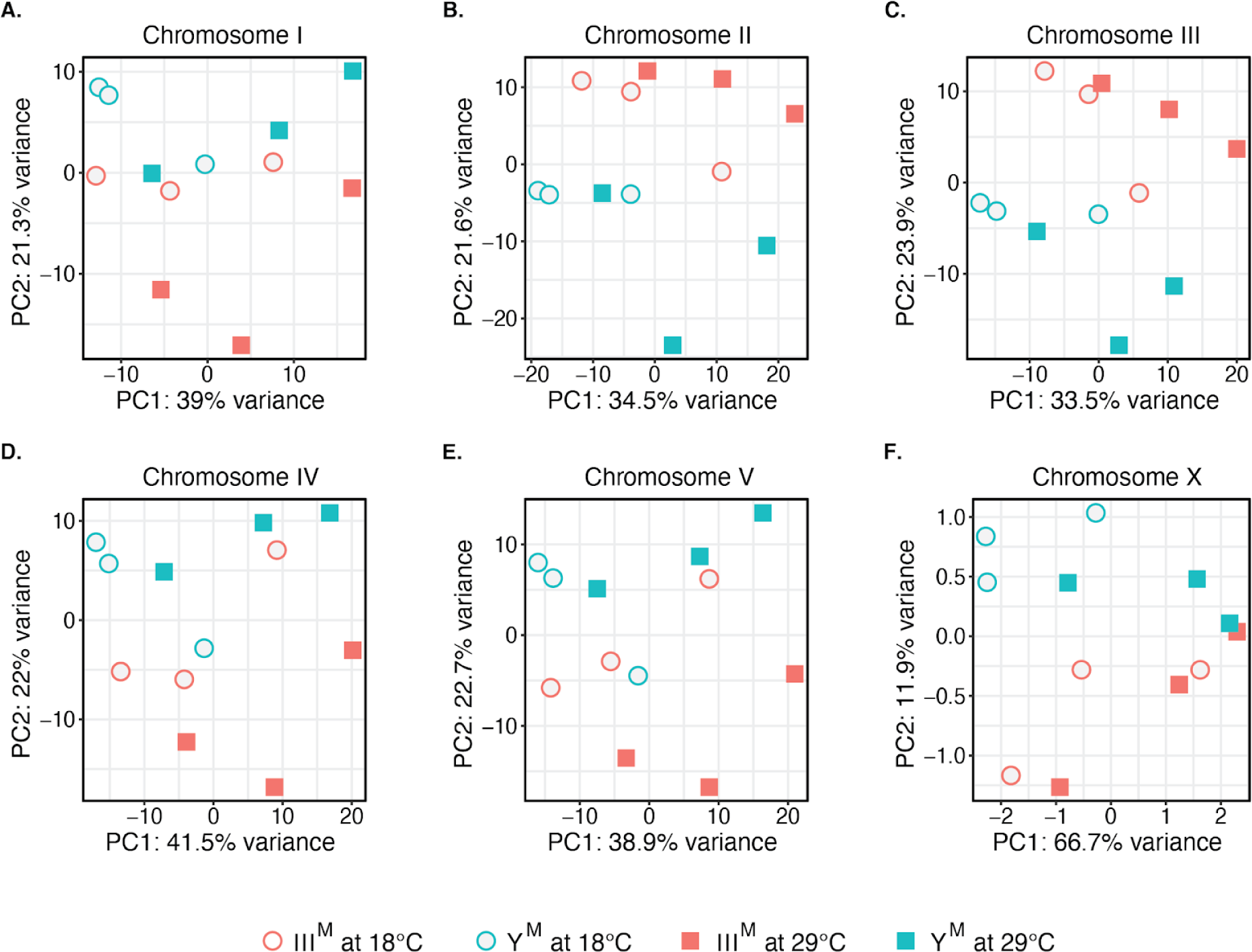
PCA plots of gene expression levels in all male heads on Chromosome I (A), Chromosome II (B), Chromosome III (C), Chromosome IV (D), Chromosome V (E), and the X Chromosome (F).

**Supplementary Figure S3.**
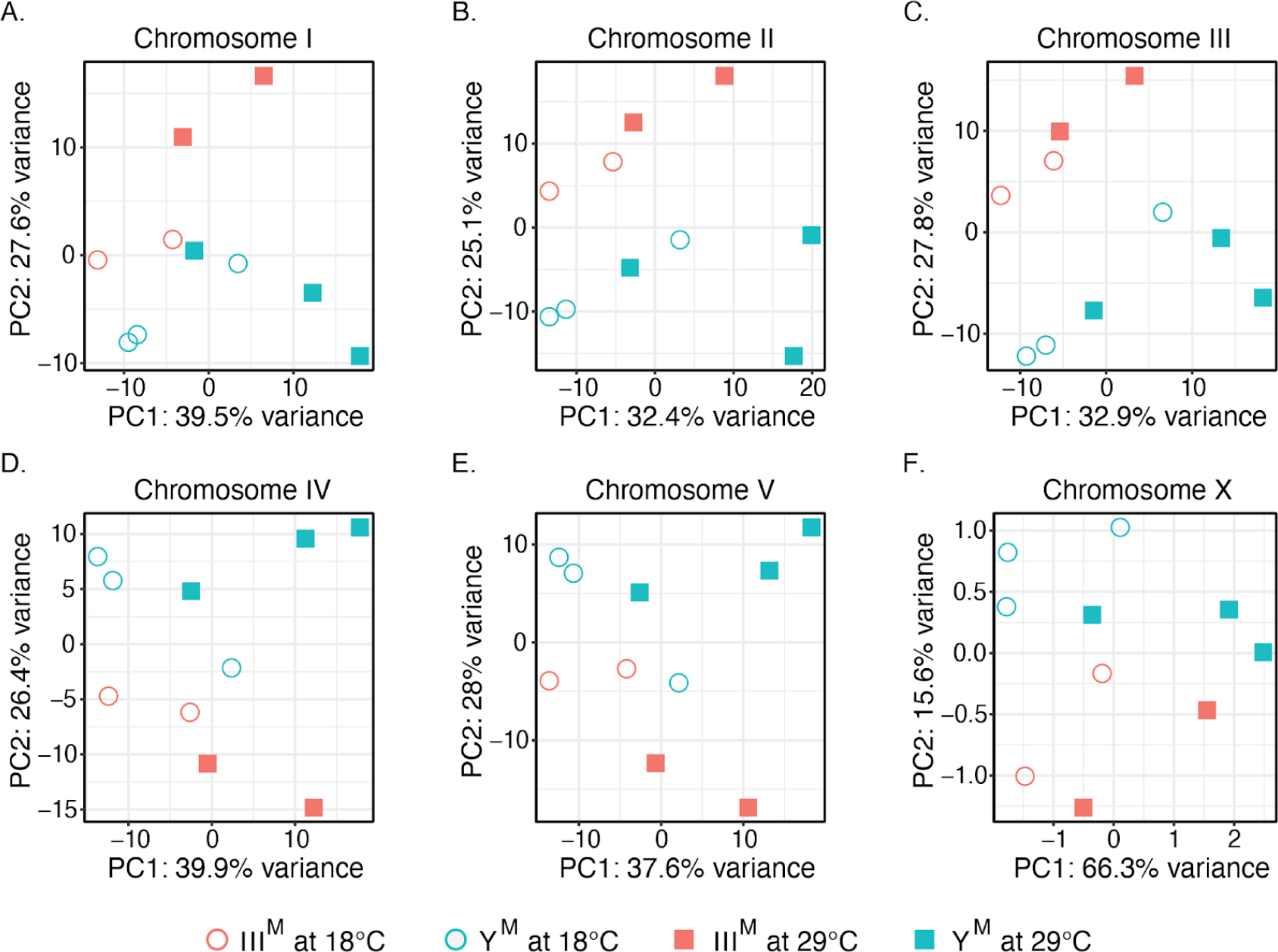
PCA plots of gene expression levels in young male heads on Chromosome I (A), Chromosome II (B), Chromosome III (C), Chromosome IV (D), Chromosome V (E), and the X Chromosome (F).

**Supplementary Figure S4.**
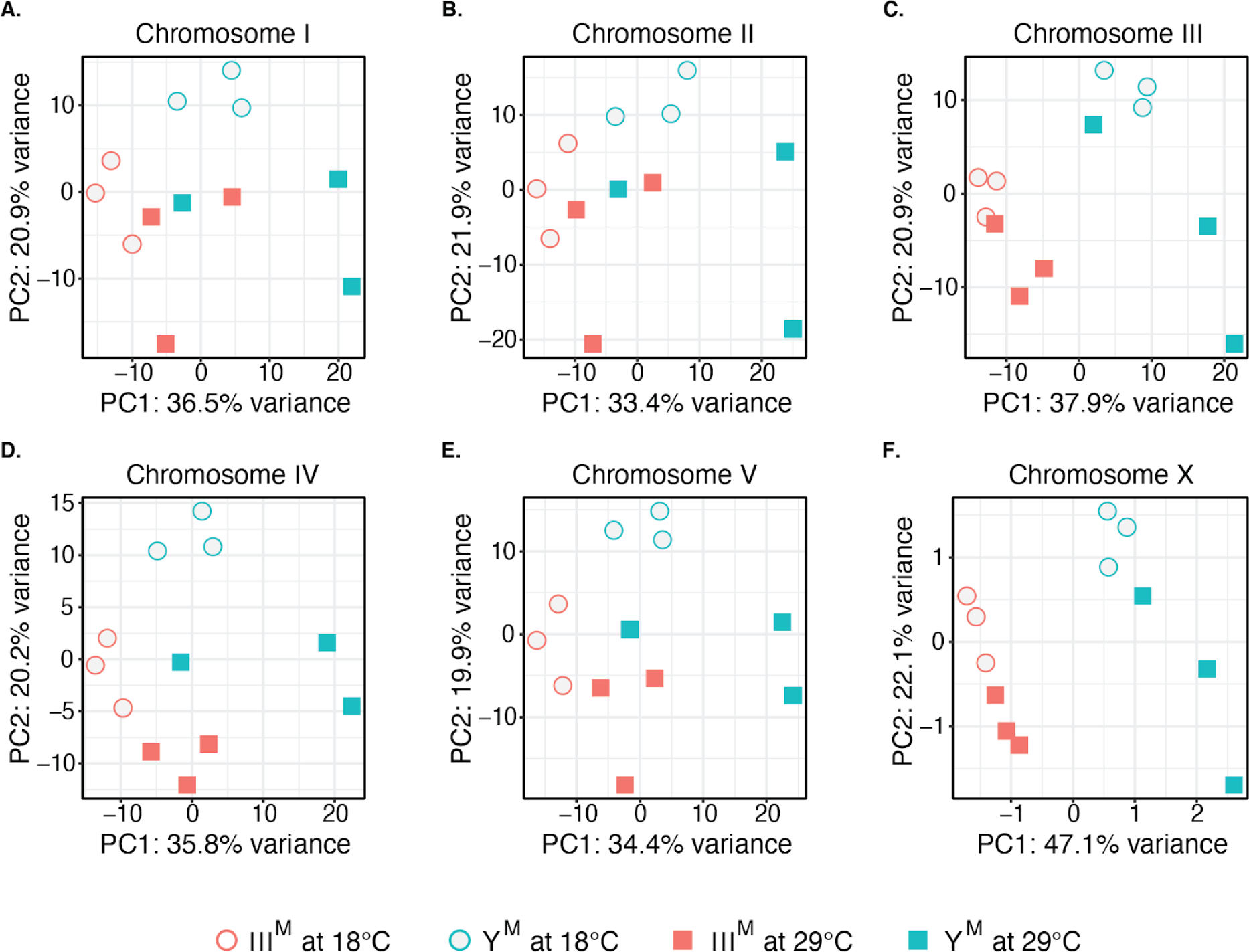
PCA plots of gene expression levels in testes on Chromosome I (A), Chromosome II (B), Chromosome III (C), Chromosome IV (D), Chromosome V (E), and the X Chromosome (F).

**Supplementary Figure S5.**
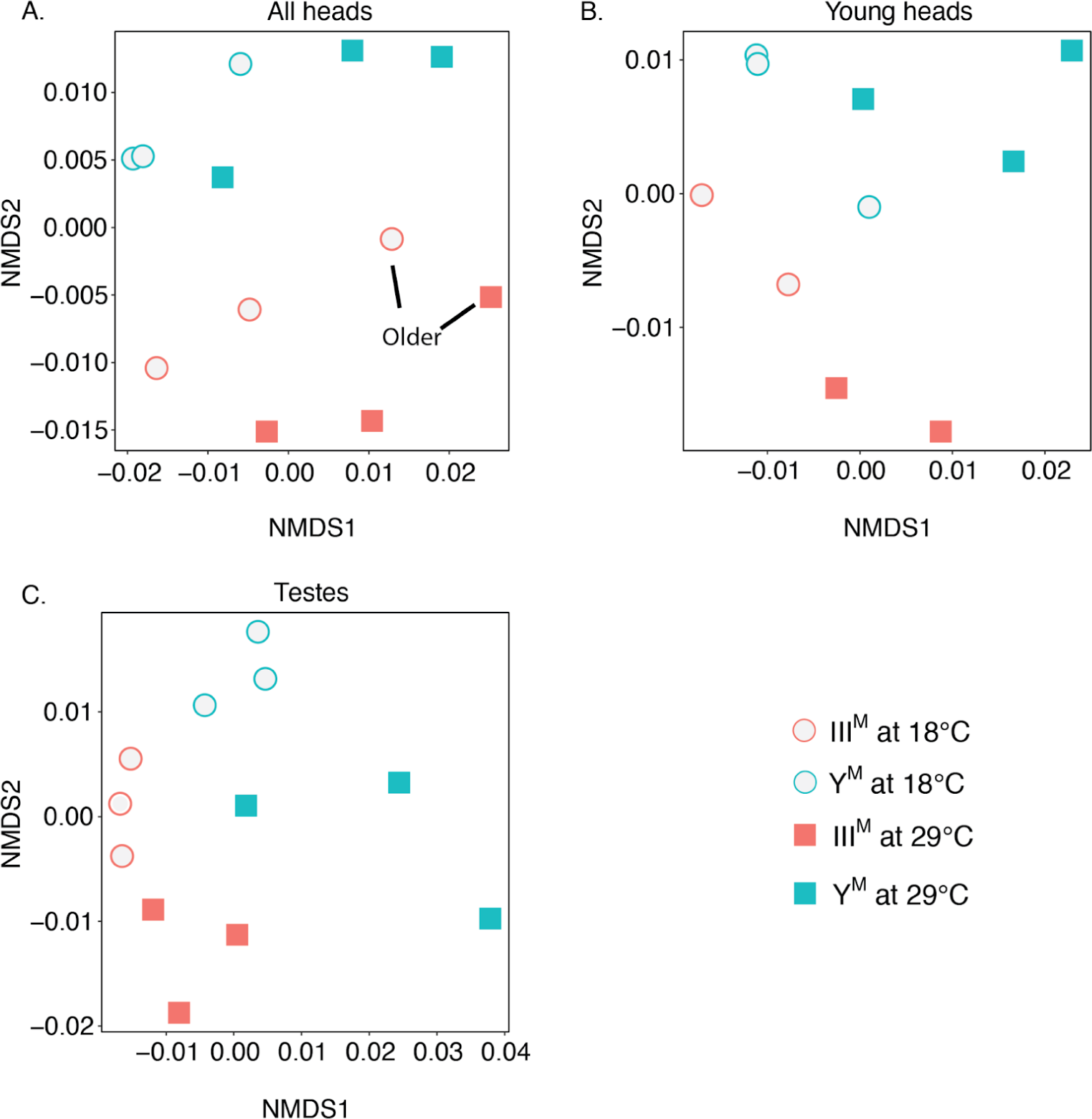
Non-metric multidimensional scaling (NMDS) plots showing gene expression profiles in all male heads (A), young male heads (B) and testes (C).

**Supplementary Figure S6.**
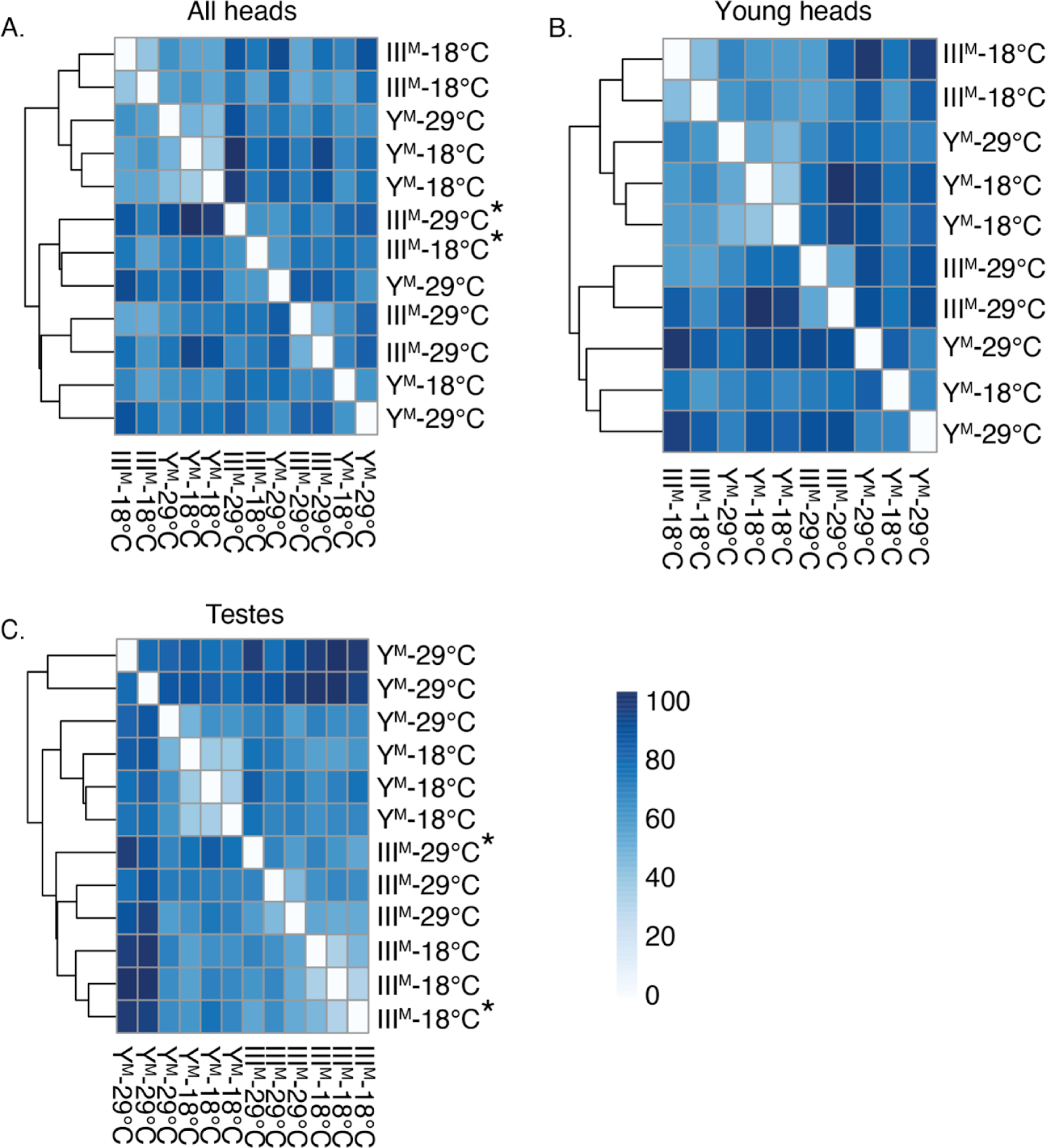
Heatmaps and dendrograms showing hierarchical clustering in all male heads (A), young male heads (B), and testes (C). Samples were compared using the Euclidean distance between regularized log transformed read counts. Color (0-80) refers to the euclidean distance between samples. Asterisks indicate older samples.

**Supplementary Figure S7:**
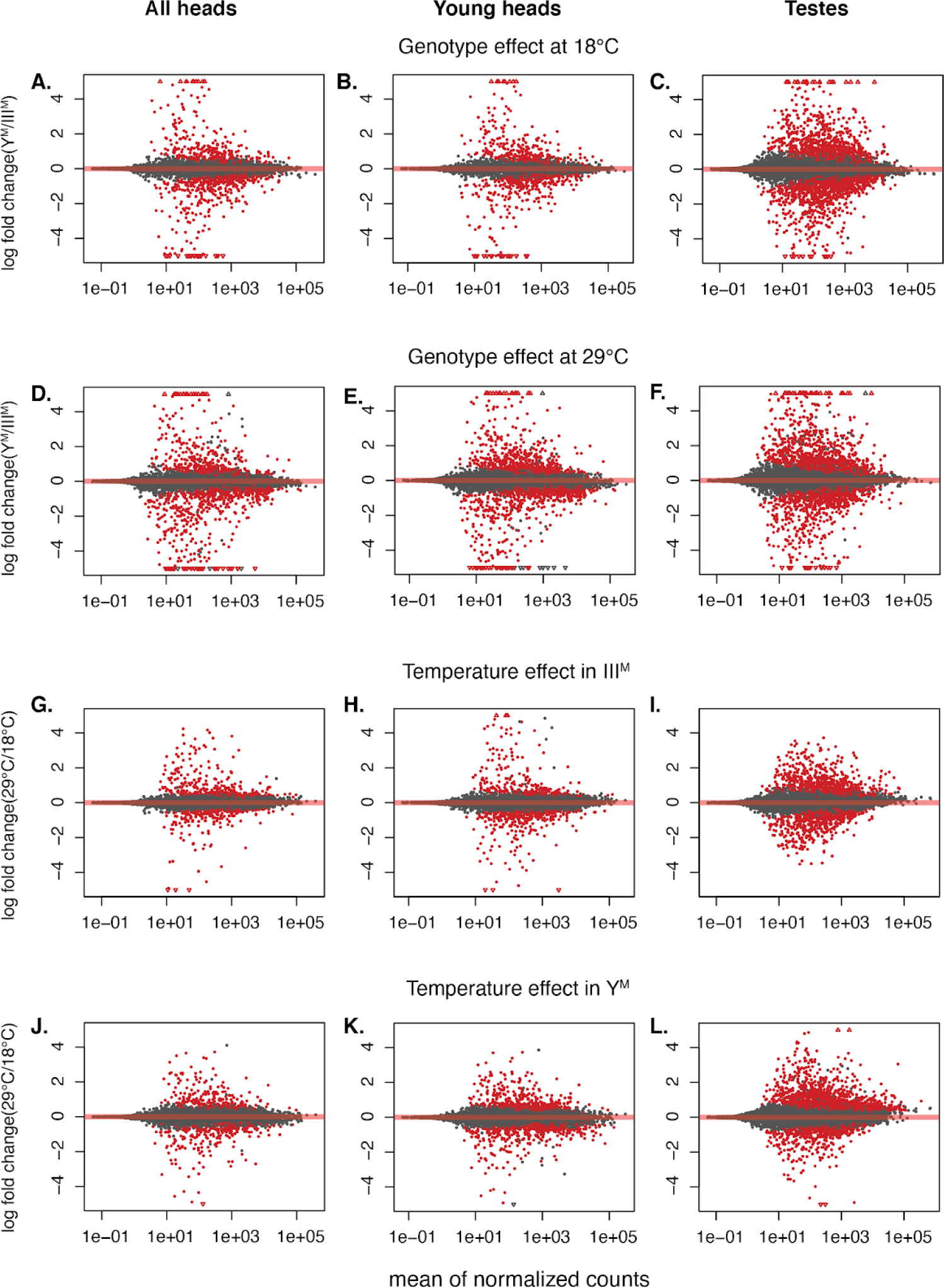
MA plots showing differential expression of genes because of: genotype effect at 18°C in all male heads (A), young male heads (B), or testes (C); genotype effect at 29°C in all male heads (D), young male heads (E), or testes (F); temperature effect in III^M^ males in all male heads (G), young male heads (H), or testes (I); and temperature effect in Y^M^ males in all male heads (J), young male heads (K), or testes (L). The log fold change values were shrunk using the lfcShrink() function in DESeq2 with the adaptive shrinkage estimator (Stephens 2016). Each dot represents a gene. Red dots represent genes that are significantly differentially expressed (p<0.05).

**Supplementary Figure S8.**
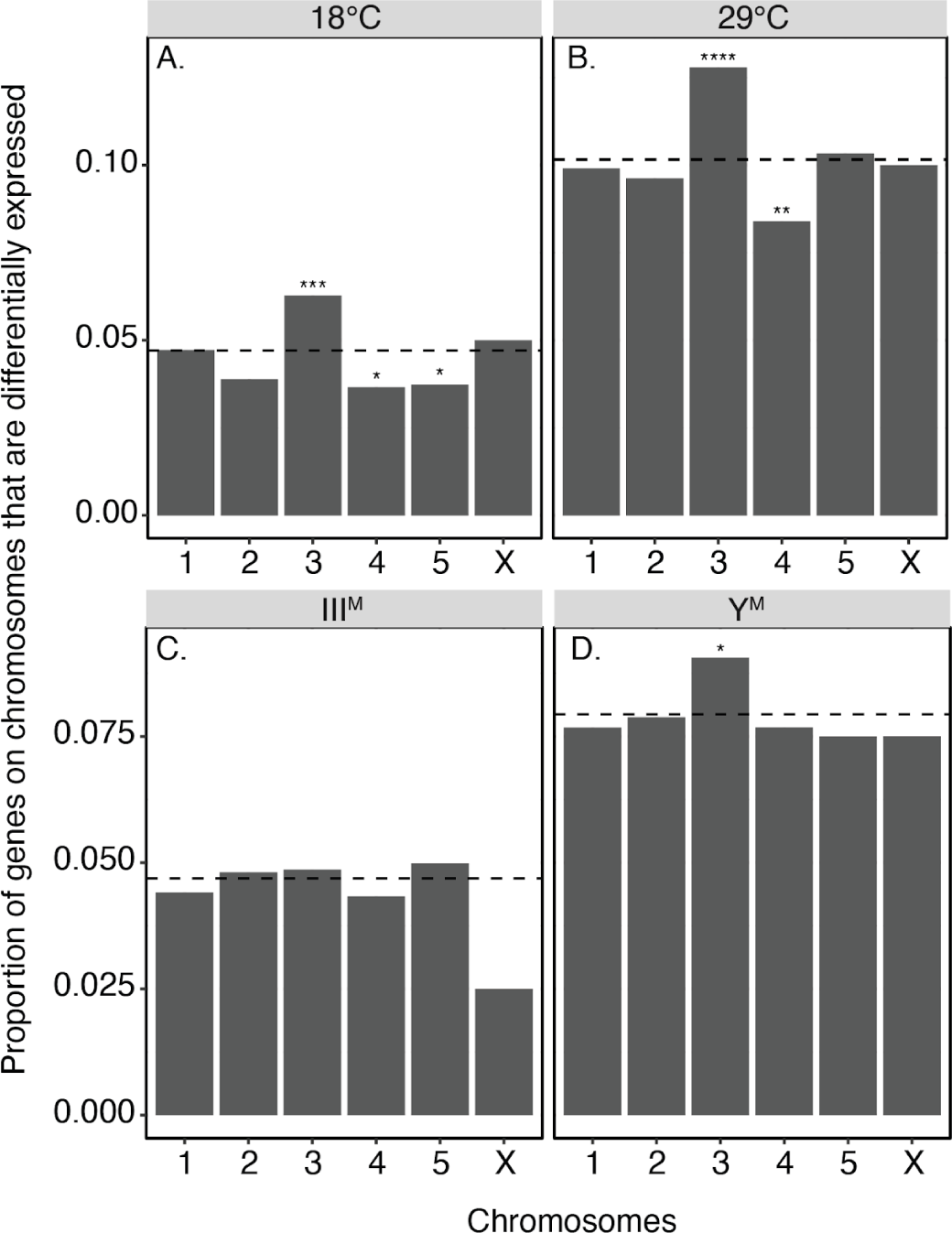
Proportion of genes that are differentially expressed in head on each chromosome when the two older samples were excluded. Comparisons are between genotypes at 18°C (A) and 29°C (B), as well as between temperatures in III^M^ males (C) and Y^M^ males (D). Each bar represents the proportion of differentially expressed genes on a chromosome, and dashed lines show the genome-wide average. Asterisks indicate *P* values obtained from Fisher’s exact test comparing the number of differentially expressed genes on a chromosome, the number of non-differentially expressed genes on a chromosome, and the number of differentially and non-differentially expressed genes across all other chromosomes, after Bonferroni correction (\**P* < 0.05, \*\**P* < 0.005, \*\*\**P* < 0.0005, \*\*\*\**P* < 0.00005, \*\*\*\*\**P* < 0.000005). When we exclude the two older III^M^ head samples, there is a modest enrichment of third chromosome genes that are significantly differentially expressed between temperatures in Y^M^ heads (Fisher’s Exact Test, Odds ratio = 1.2, 95%CI: 1.01-1.42, *P* = 0.03).

**Supplementary Figure S9:**
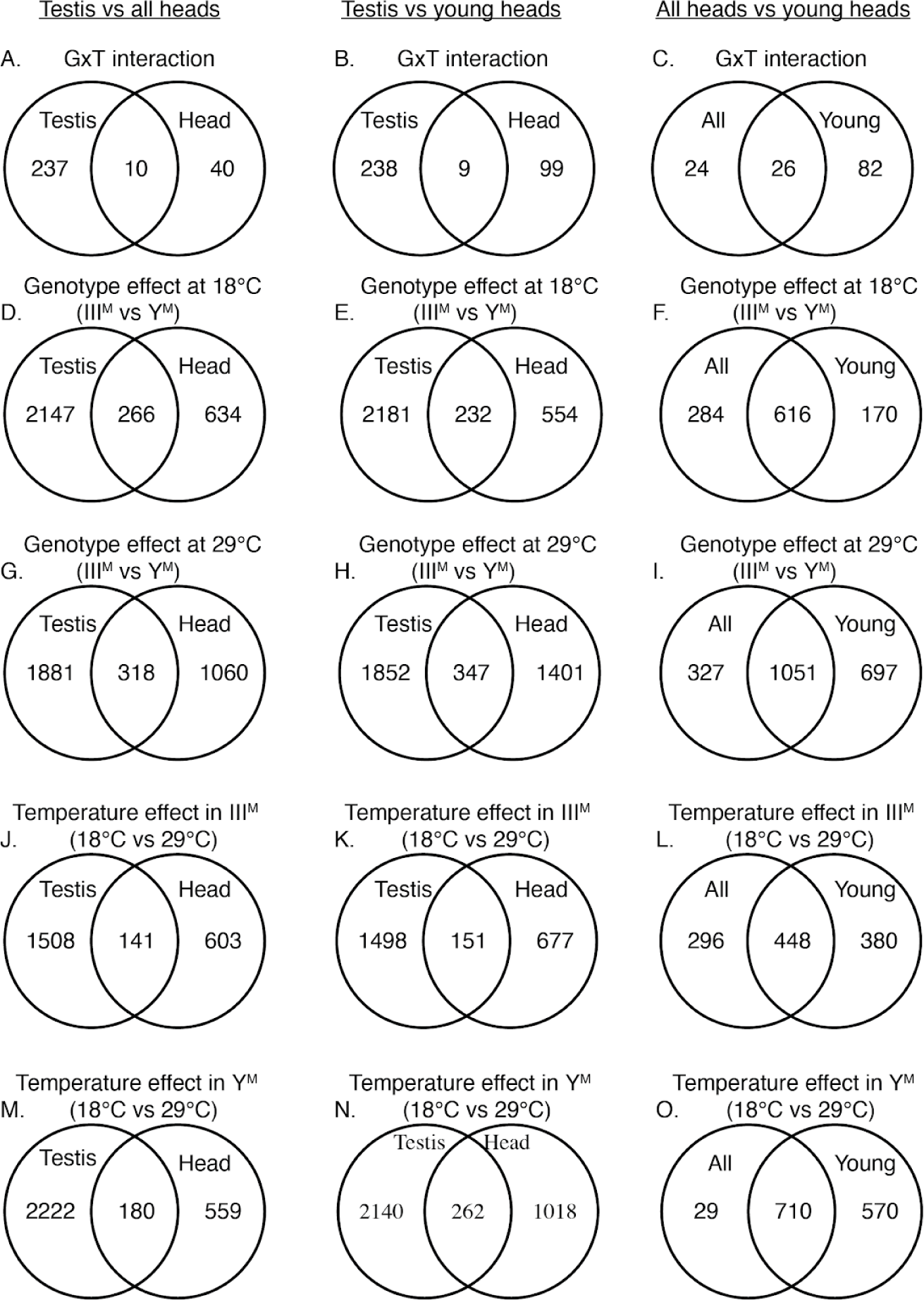
Venn diagrams show the number of significantly differentially expressed genes in testes vs all male heads (first column), testes vs young male heads (second column), and all male heads vs young male heads (third column). There is an excess of genes in the overlapping portion in all comparisons (*P*<0.05 using a *z*-test of proportions).

**Supplementary Figure S10.**
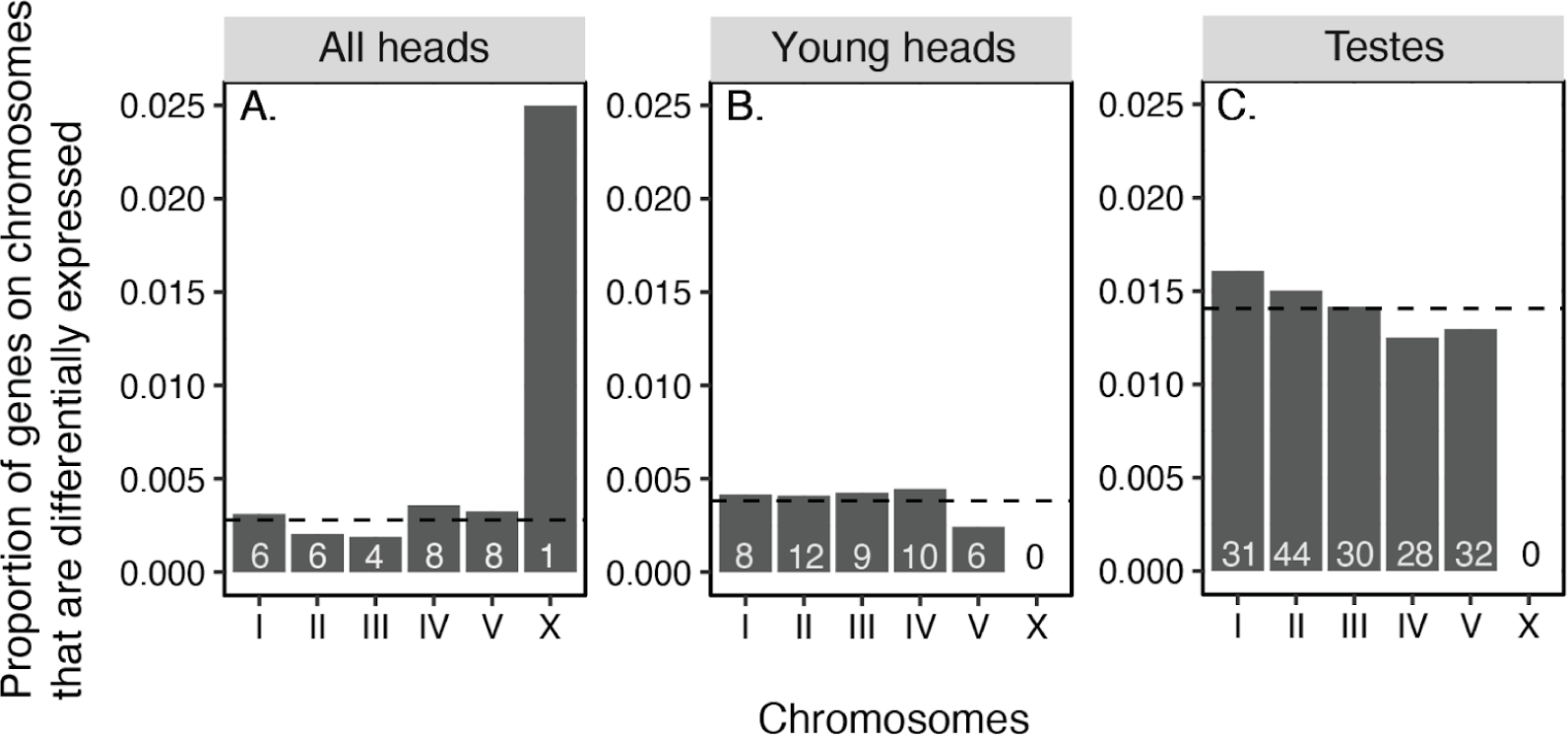
No chromosomes are enriched for significantly differentially expressed genes as a result of G×T interactions in all male heads (A), young male heads (B), or testes (C). The dashed line represents the genome-wide proportion of significantly differentially expressed genes. The numbers within each bar represent the number of significantly differentially expressed genes on that chromosome. Not all genes with significant G×T effects on expression are assigned to chromosomes, and only genes assigned to chromosomes are plotted. The X chromosome has <100 genes, which is much less than the other chromosomes which have 1000s of genes. Therefore a single G×T interaction on the X chromosome appears as a large proportion of genes. However, the small total number of X chromosome genes means this large proportion is not a significant excess.

**Supplementary Figure S11.**
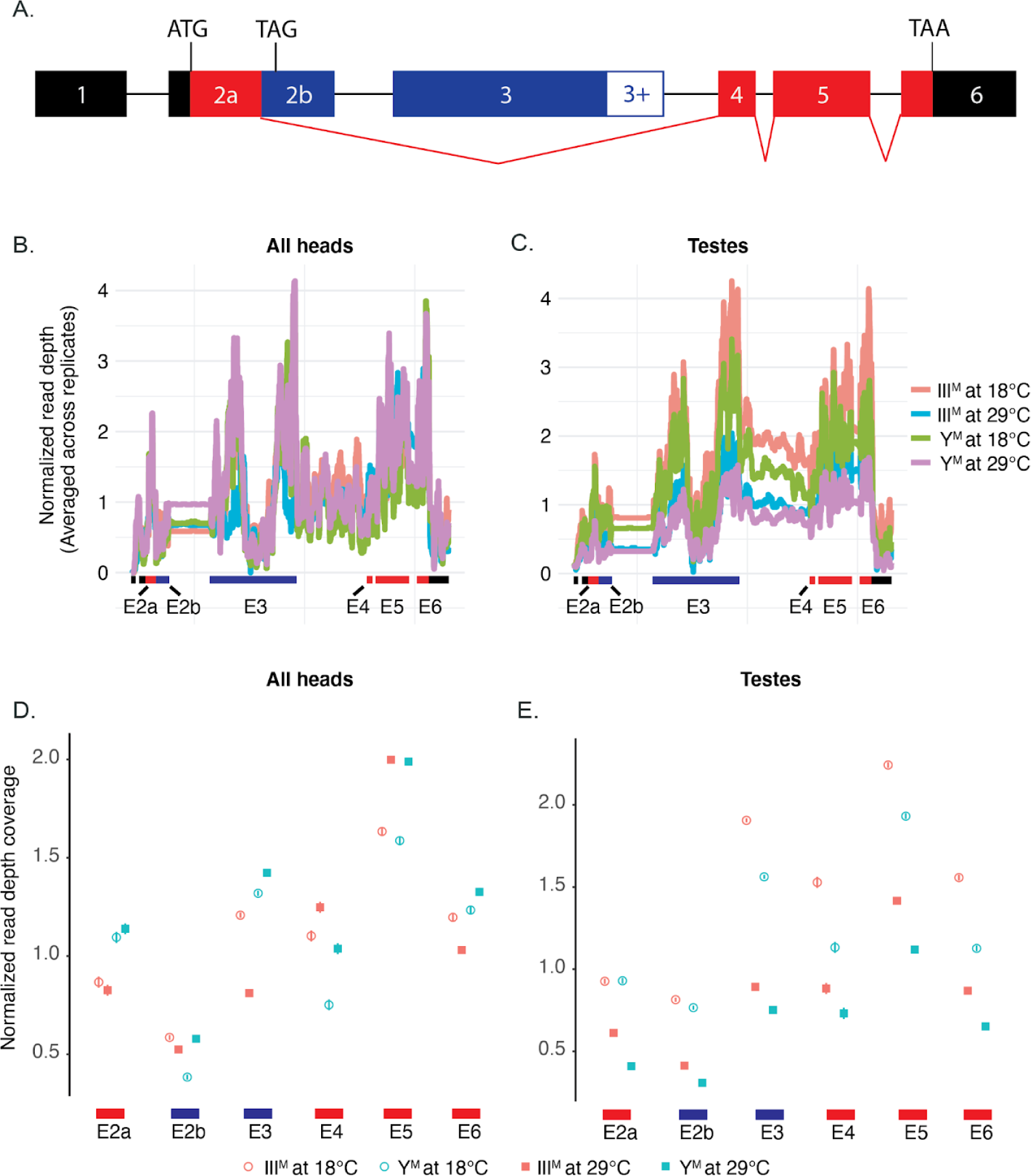
(A) Schematic representation of the *Md-tra* locus based on DNA sequencing, cDNA clones, and RNA-Seq data (Hediger *et al*. 2010; Scott *et al*. 2014). Splicing of the female-determining transcript is illustrated by the red diagonal lines connecting exons, and exons that contain protein-coding sequence of the female-determining splice variant are in red. Exons found in the male isoforms are shown in blue. The start and stop codon locations are shown. The expression of *Md-tra* across all exons is shown for RNA-seq data collected from all male heads (B and D) and testis (C and E). Error bars represent standard error (most standard error estimates are smaller than the size of the points, and thus cannot be seen in the graph).

**Supplementary Figure S12.**
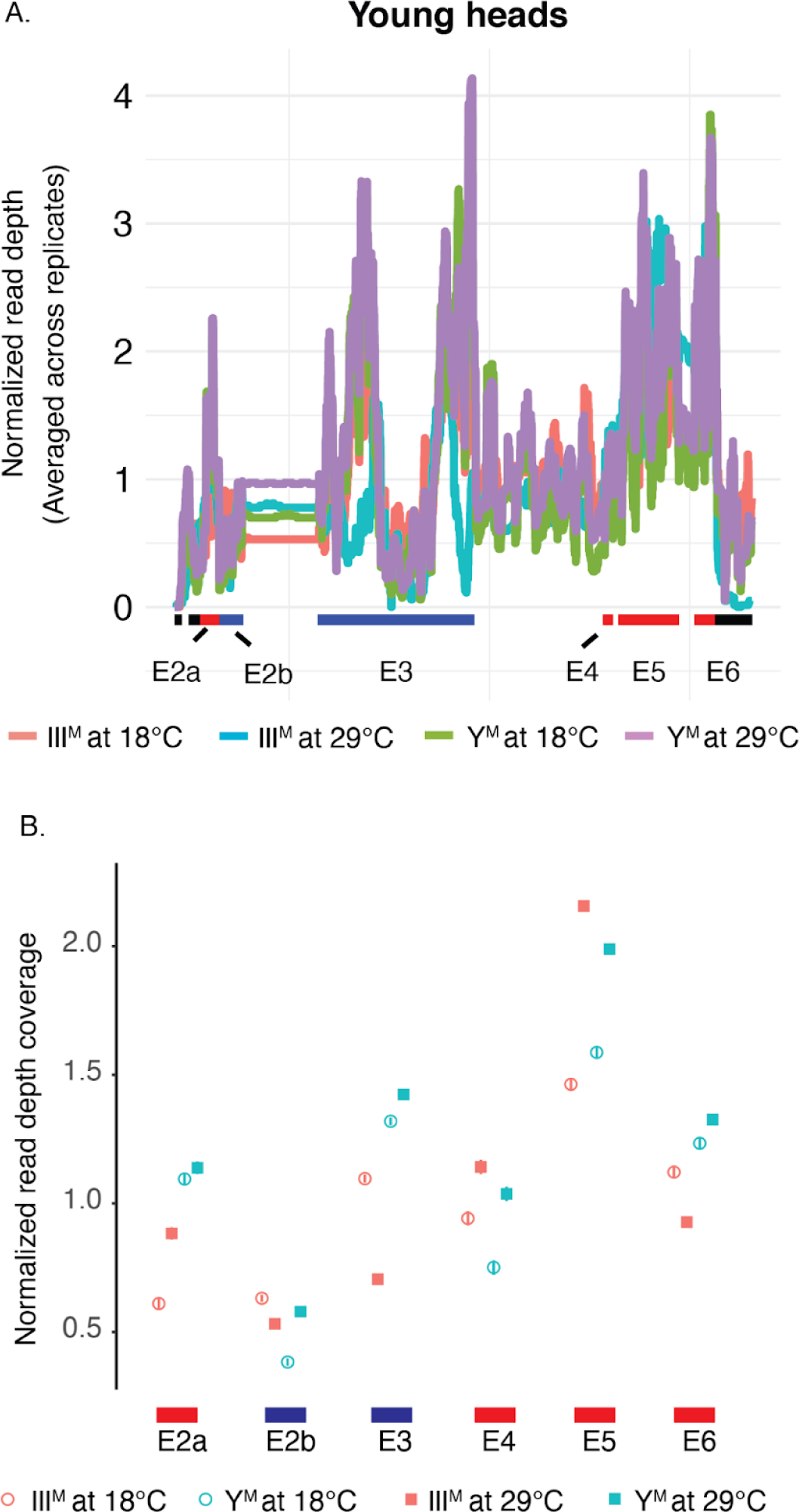
Expression of *Md-tra* across all exons is shown for RNA-seq data collected from young male heads. Error bars represent standard error (most standard error estimates are smaller than the size of the points, and thus cannot be seen in the graph). Exons that contain protein-coding sequence of the female-determining splice variant (E2a, E4, E5,and E6) are in red. Exons found in the male isoforms (E2b and E3) are shown in blue.

**Supplementary Figure S13.**
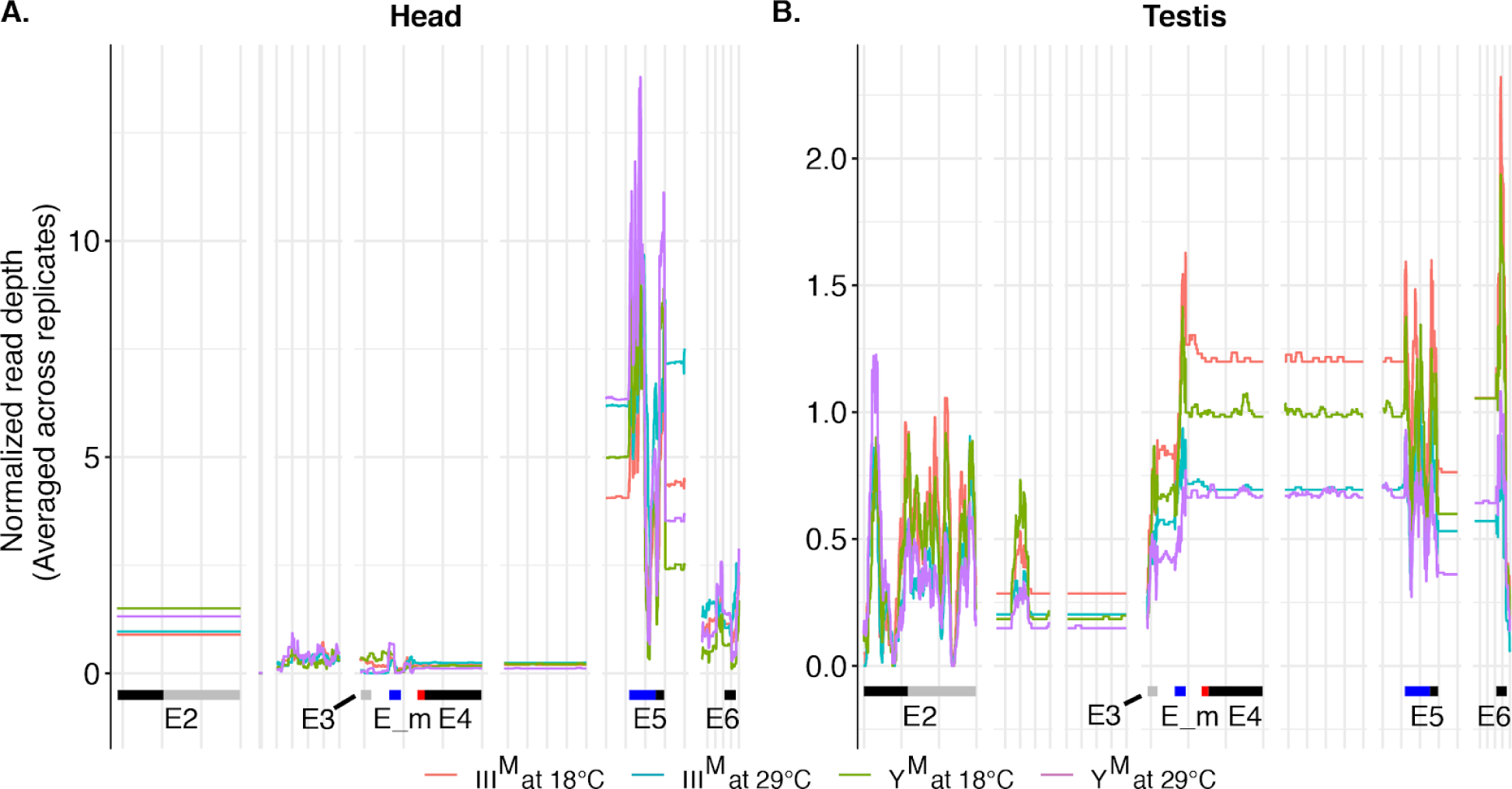
Read depth coverage of *Md-dsx* in all male heads (A) and testis (B) in each G×T combination. Exons are shown along the x-axis. Exons in the male-determining isoform are shown in blue, and exons in the female-determining isoform are shown in red. Expression of *Md-dsx* is not significantly affected by G×T interactions in either head or testis (Supplementary Tables 2 and 3). Usage of male-specific exon E_m (ANOVA, *P* = 0.0004), exon E5 (ANOVA, *P* = 0.013), and the female specific exon E4 (ANOVA, *P* < 2.2e-16) are affected by G×T interactions in testis. However, these G×T interactions are not in the directions expected if mis-splicing of *Md-dsx* is responsible for maintaining Y^M^-III^M^ clines. In head, usage of E_m (ANOVA, *P* < 2.2 e-16) and Exon 4 (ANOVA, *P* < 2.2 e-16) is affected by G×T interactions. Similar to testis, effects of the G×T interactions in head are also not in the directions expected if mis-splicing of *Md-dsx* is responsible for maintaining Y^M^-III^M^ clines.

**Supplementary Figure S14.**
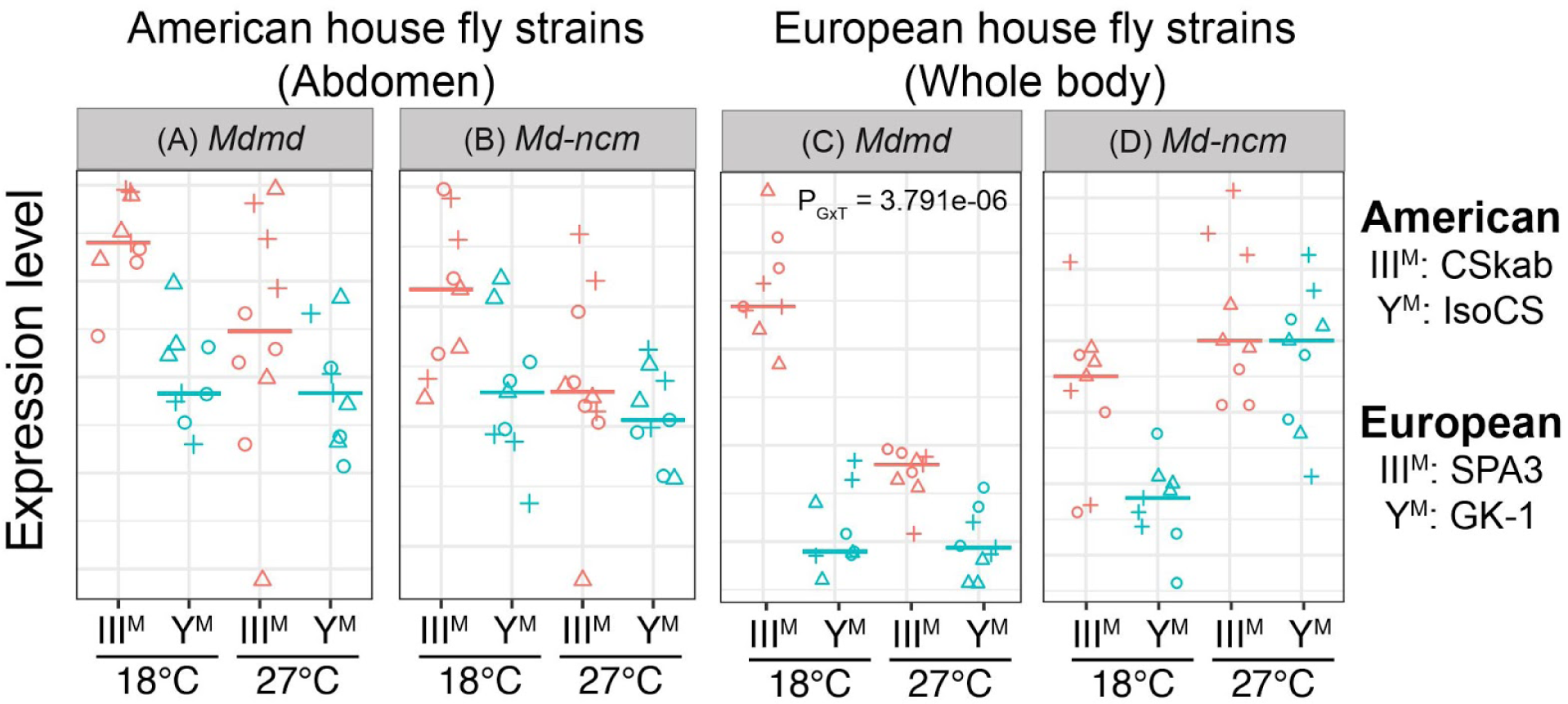
Expression levels of *Mdmd* (A), and *Md-ncm* (B) in the abdomens of CSkab (III^M^) and IsoCS (Y^M^), and expression levels of *Mdmd* (C), and *Md-ncm* (D) in the whole body of SPA3 (III^M^) and GK-1 (Y^M^) males raised at 18°C and 27°C. Each data point is a technical replicate that has been normalized by dividing by the control gene within that replicate, and points with the same shape are from the same biological replicate. The horizontal line indicates the median across all replicates.

**Supplementary Figure S15.**
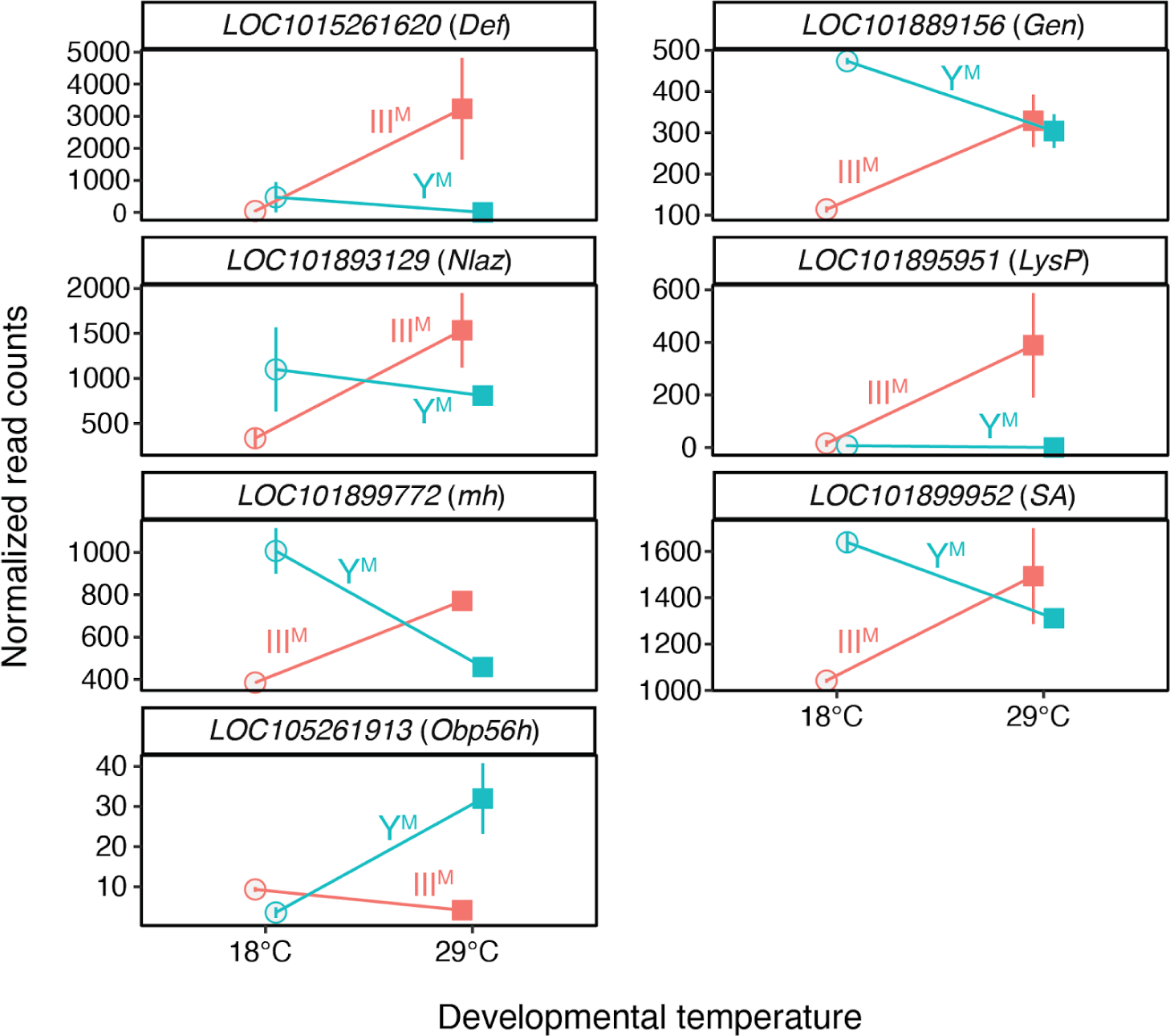
Normalized read counts in young male heads (excluding two older head samples) for genes that are differentially expressed and have consistent expression patterns with the analysis when the two older head samples are included (Figure 3A). *Nlaz* is not significantly differentially expressed when only younger heads are considered (*P* =0.08), however the pattern is consistent with the analysis of all heads. All other genes are significantly differentially expressed. Error bars represent standard error of mean.

## Supplementary Tables

**Supplementary Table S1.**
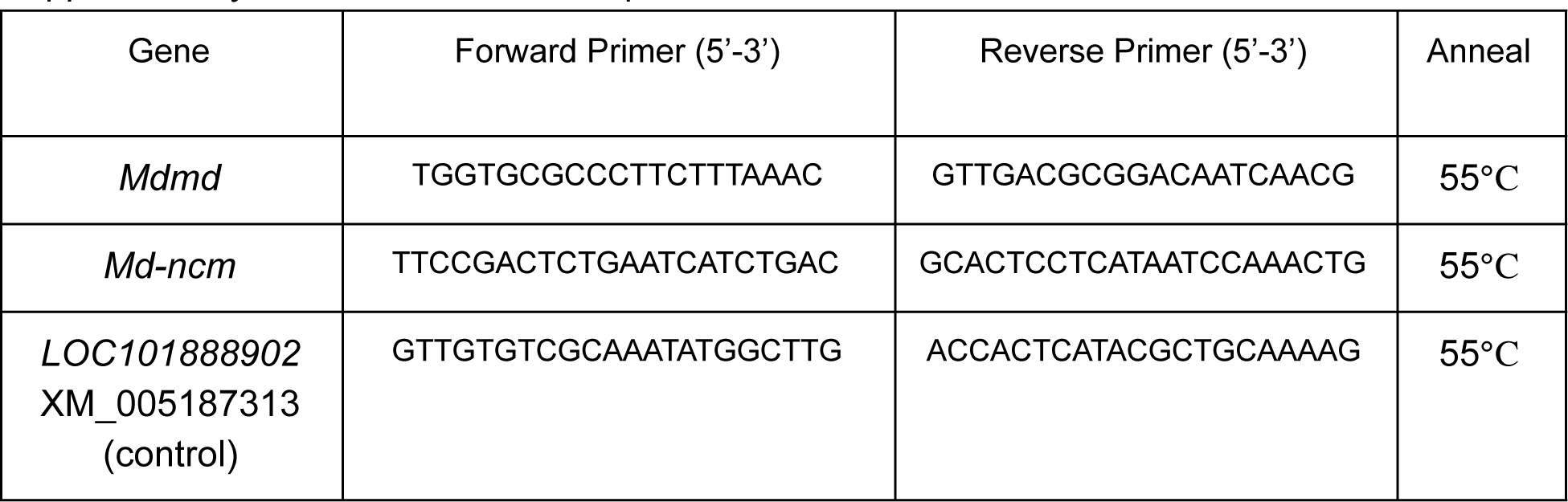
Primers for qPCR

**Supplementary Table S2:** RNA-seq reads from head mapped to the reference genome

| Sample | Geno<br>-type | Mapped<br>reads | Total reads | % mapped<br>reads | Uniquely<br>mapped reads |
| --- | --- | --- | --- | --- | --- |
| CSrab_18C_head1 | III<sup>M</sup> | 23,386,737 | 30,872,584 | 75.75 | 18,211,108 |
| CSrab_18C_head2 | III<sup>M</sup> | 36,498,365 | 48,036,045 | 75.98 | 28,405,569 |
| CSrab_18C_head3 * | III<sup>M</sup> | 25,899,682 | 33,922,170 | 76.35 | 20,123,259 |
| CSrab_29C_head1 | III<sup>M</sup> | 29,078,715 | 35,935,602 | 80.92 | 24,626,862 |
| CSrab_29C_head2 | III<sup>M</sup> | 32,542,292 | 40,966,941 | 79.44 | 26,979,708 |
| CSrab_29C_head3 * | III<sup>M</sup> | 45,976,216 | 60,550,809 | 75.93 | 35,694,337 |
| IsoCS_18C_head1 | Y<sup>M</sup> | 23,338,684 | 30,429,230 | 76.70 | 18,470,903 |
| IsoCS_18C_head2 | Y<sup>M</sup> | 23,593,266 | 30,809,521 | 76.58 | 18,657,918 |
| IsoCS_18C_head3 | Y<sup>M</sup> | 20,170,741 | 26,500,047 | 76.12 | 15,690,817 |
| IsoCS_29C_head1 | Y<sup>M</sup> | 30,777,222 | 40,045,778 | 76.86 | 24,278,617 |
| IsoCS_29C_head2 | Y<sup>M</sup> | 36,530,758 | 47,069,689 | 77.61 | 28,848,124 |
| IsoCS_29C_head3 | Y<sup>M</sup> | 29,711,982 | 38,668,539 | 76.84 | 23,352,024 |

Asterisks indicate the two older sample

**Supplementary Table S3:** RNA-seq reads from testis mapped to the reference genome

| Sample | Geno<br>-type | Mapped<br>reads | Total reads | % mapped<br>reads | Uniquely<br>mapped reads |
| --- | --- | --- | --- | --- | --- |
| CSrab_18C_testis1 | III<sup>M</sup> | 18,951,438 | 24,457,263 | 77.49 | 15,651,117 |
| CSrab_18C_testis2 | III<sup>M</sup> | 18,154,085 | 23,758,773 | 76.41 | 14,916,846 |
| CSrab_18C_testis3 | III<sup>M</sup> | 24,260,793 | 32,425,899 | 74.82 | 19,961,330 |
| CSrab_29C_testis1 | III<sup>M</sup> | 34,972,612 | 46,278,674 | 75.57 | 28,989,480 |
| CSrab_29C_testis2 | III<sup>M</sup> | 25,449,937 | 32,944,157 | 77.25 | 21,232,767 |
| CSrab_29C_testis3 | III<sup>M</sup> | 38,081,673 | 51,434,420 | 74.04 | 31,713,175 |
| IsoCS_18C_testis1 | Y<sup>M</sup> | 39,350,279 | 49,827,408 | 78.97 | 32,970,057 |
| IsoCS_18C_testis2 | Y<sup>M</sup> | 24,820,579 | 31,850,279 | 77.93 | 20,609,110 |
| IsoCS_18C_testis3 | Y<sup>M</sup> | 31,389,010 | 39,651,139 | 79.16 | 26,217,670 |
| IsoCS_29C_testis1 | Y<sup>M</sup> | 48,781,311 | 65,167,161 | 74.86 | 40,005,841 |
| IsoCS_29C_testis2 | Y<sup>M</sup> | 30,454,185 | 41,083,650 | 74.13 | 24,992,406 |
| IsoCS_29C_testis3 | Y<sup>M</sup> | 38,393,746 | 48,460,189 | 79.23 | 32,299,686 |

Supplementary Table S4. DESeq2 result for all male heads. Data provided in a separate file.

Supplementary Table S5. DESeq2 result for young male heads. Data provided in a separate file.

Supplementary Table S6. DESeq2 result for testes. Data provided in a separate file.

## Notes

https://www.ncbi.nlm.nih.gov/geo/query/acc.cgi?acc=GSE136188

### Competing Interest Statement

The authors have declared no competing interest.

### Summary of Updates

This version of the manuscript has been revised in response to reviewer comments after submission to a journal.

